# Protein S-glutathionylation confers cell resistance to ferroptosis

**DOI:** 10.1101/2024.05.03.592374

**Authors:** Yi Ju, Yuting Zhang, Yiming Qiao, Xiaolin Tian, Yufan Zheng, Tao Yang, Baolin Niu, Xiaoyun Li, Liu Yu, Zhuolin Liu, Yixuan Wu, Yang Zhi, Yinuo Dong, Qingling Xu, Xuening Wang, Xiaokai Wang, Yimin Mao, Xiaobo Li

## Abstract

Ferroptosis is a type of cell death that is strongly associated with the cellular redox state. Glutathione is the key to buffering lipid peroxidation in ferroptosis and can also modify proteins by S-glutathionylation under oxidative stress. Here, we showed that the strong associations among glutathione pools, protein S-glutathionylation, and susceptibility to ferroptosis existed broadly in ferroptosis induced by erastin or acetaminophen. Deficiency of CHAC1, a glutathione-degrading enzyme, led to decreased glutathione pools and reduced protein S-glutathionylation, improved liver function and attenuated hepatocyte ferroptosis upon acetaminophen challenge, which could be retarded by CHAC1 overexpression. We conducted quantitative redox proteomics in primary mouse hepatocytes to identify glutathione pool-sensitive S-glutathionylated proteins and found that S-glutathionylation is required to maintain the function of ADP-ribosylation factor 6 (ARF6). Our data suggest that aberrant ARF6 S-glutathionylation increases the labile iron pool by delaying the recycling of transferrin receptors, thereby promoting ferroptosis. Our study reveals the importance of protein S-glutathionylation in conferring cell resistance to ferroptosis.

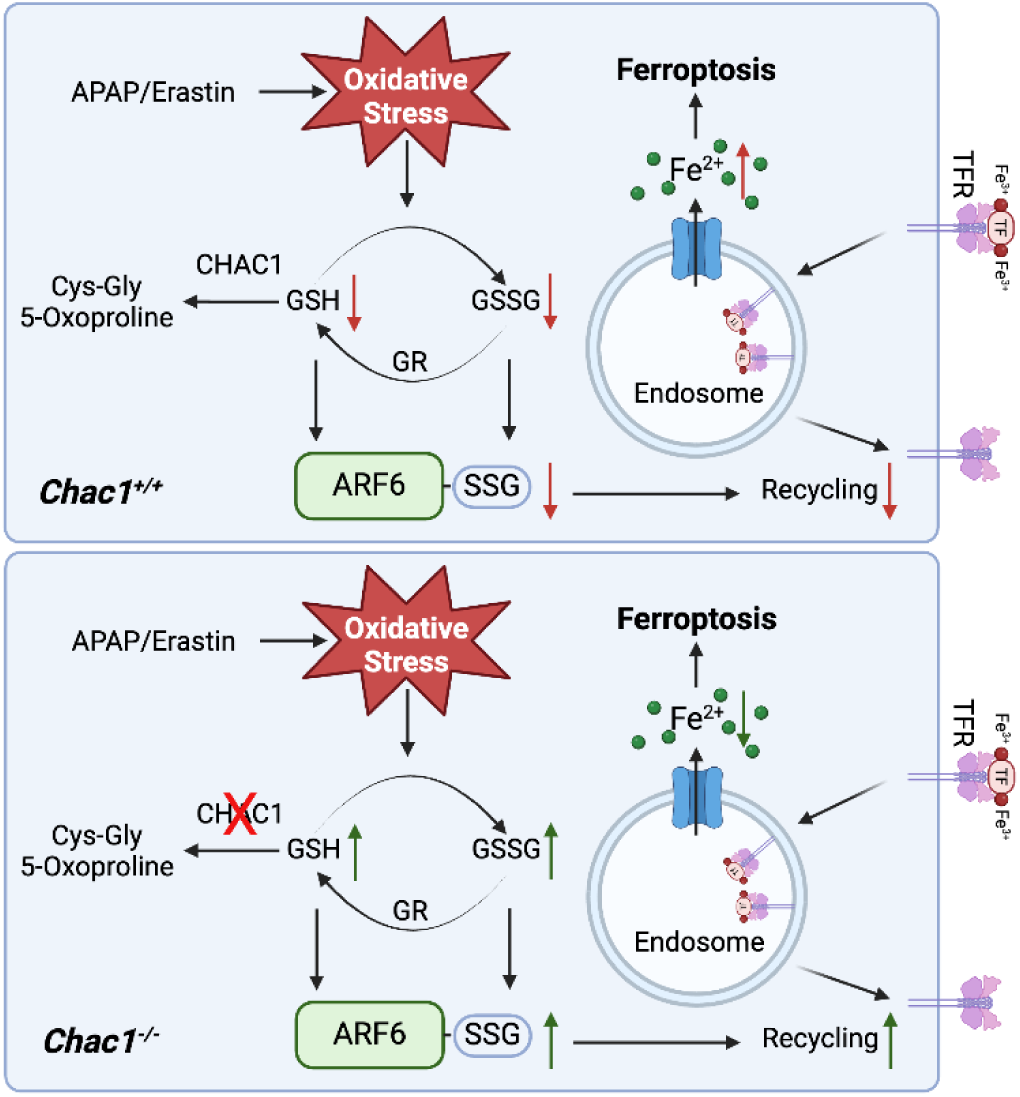

**HIGHLIGHTS:**

1. Highly upregulated CHAC1 decreases glutathione pools and protein S-glutathionylation.
2. Reduced protein S-glutathionylation associated with decreased glutathione pools promotes ferroptosis.
3. S-glutathionylation of ARF6 at Cys90 promotes ARF6 activation.
4. Reduced S-glutathionylation of ARF6 provides a labile iron pool to drive ferroptosis.

## INTRODUCTION

Ferroptosis, the term first coined in 2012, is an iron-dependent form of cell death driven by lipid peroxidation. Ferroptosis is implicated in processes, such as aging, development, iron-overload diseases, organ injury, neurodegeneration, infectious diseases, autoimmune diseases, tumorigenesis (Stockwell, 2022). Ferroptosis inducers have the potential to be used in cancer treatment, whereas ferroptosis inhibitors have the potential to treat organ injury and neurodegeneration(Liang, Minikes et al., 2022). The solute carrier family 7 member 11 (SLC7A11) -glutathione (GSH) axis is an essential surveillance mechanism for defence against ferroptosis, in which GSH acts as a cofactor for the selenoenzyme glutathione peroxidase 4 (GPX4) to scavenge lipid peroxide (Jiang, Stockwell et al., 2021). GSH, the most abundant antioxidant in cells, ranges from 1 to 10 mM (Labarrere & Kassab, 2022). Upon reduction of lipid peroxide levels, GSH is converted to oxidised glutathione (GSSG), which can be reversed by glutathione reductase and the cofactor NADPH. Certain biological processes maintain GSH homeostasis and resolve cellular redox reactions. GSH homeostasis plays a crucial role in several pathological processes (Chai & Mieyal, 2023).

Glutathione can also modify proteins via S-glutathionylation (protein-SSG). Under conditions of oxidative stress, cysteine residues in proteins can be oxidised to reversible sulfenic acid, sulfonic acid, and ultimately to irreversible sulfonic acid by reactive oxygen species; they can also be oxidised to reversible S-nitrosothiols by reactive nitrogen species (Matsui, Ferran et al., 2020). Irreversible oxidative posttranslational modifications (OPTM) of cysteine residues can permanently damage or degrade proteins. Protein S-glutathionylation is a reversible and dynamic OPTM that forms a disulfide bond between glutathione and a cysteine residue in the target protein. Protein-SSG can occur spontaneously without enzymes. Under oxidative stress, the thiol groups of proteins are first oxidised into such high-oxidation groups as sulfenic acid and S-nitrosothiols, which then react with GSH to form S-glutathionylation, or GSSG can directly modify the proteins. Protein-SSG could be reversed to protein-SH upon resolution of oxidative stress (Mailloux, 2020, Musaogullari & Chai, 2020). Protein-SSG can protect cysteine residues from irreversible oxidation under oxidative stress or regulates enzyme activity, localisation, protein interaction, and stability, and affects the occurrence and development of diseases, including aging, neurodegeneration, liver diseases, cardiovascular diseases, inflammation, and tumours (A, Diwakar et al., 2022, Checconi, Limongi et al., 2019, Musaogullari & Chai, 2020, Rashdan, Shrestha et al., 2020). Despite the potential association between protein-SSG and the redox state, as well as the significance of redox imbalance in ferroptosis, to the best of our knowledge, no study has focused on the role of protein-SSG in ferroptosis.

Emerging evidence has revealed that certain enzymes participate in the regulation of protein-SSGs. Glutathione transferase π (GSTPπ) and LanC-like protein (LanCL) promote SSG formation, while GLRX and glutathione transferase omega (GSTO) catalyse deglutathionylation (Oppong, Schiff et al., 2023). GLRX is the most well-studied catalytic enzyme involved in deglutathionylation. Researchers have studied target protein S-glutathionylation by manipulating GLRX. Nevertheless, protein S-glutathionylation is strongly associated with glutathione availability. Both GSH and GSSG are substrates for protein S-glutathionylation under oxidative stress. However, the important role of glutathione pools in protein S-glutathionylation is unknown, especially in the context of oxidative stress-related ferroptosis.

The maintenance of GSH homeostasis requires the coordination of GSH biosynthesis, transport, efflux, peroxidases (GPX4), reductases, consumption, and degradation (Bachhawat & Kaur, 2017). CHAC1 is the enzyme that catalyzes the degradation of intracellular GSH. In 2009, mammalian CHAC1 was initially identified as a pro-apoptotic endoplasmic reticulum stress. Subsequently, CHAC1 was found to function as an r-glutamyl cyclotransferase, acting specifically to convert GSH to Cys-Gly and 5-oxoproline in yeast, cell-free systems, and mammalian cell models (Chi, Byrne et al., 2014, Crawford, Prescott et al., 2015, Cui, Zhou et al., 2021, Kumar, Tikoo et al., 2012, Tsunoda, Avezov et al., 2014). CHAC1 inactivation effectively preserves GSH in mouse models (Li, Lu et al., 2023). Dr. Brent R. Stockwell’s lab in Columbia University report that the gene expression of *CHAC1, PTGS2, SLC7A11*, and *ACSL4* was induced during ferroptosis, and their upregulation detected by qPCR can be regarded as an indicator of ferroptosis (Dixon, Patel et al., 2014, Stockwell, 2022). Subsequently, CHAC1 has been extensively used as a ferroptosis marker, both in vitro and in vivo.

This study found that GLRX levels were not markedly elevated compared to CHAC1 levels during ferroptosis. In several cell lines, no changes were observed in GLRX levels, whereas protein-SSG levels were altered, suggesting the importance of the glutathione pool in regulating protein-SSG levels. Therefore, we studied the role of SSG in ferroptosis by manipulating CHAC1 expression. Here, we showed that decreased protein-SSG induced by CHAC1 can promote acetaminophen (APAP)- or erastin-induced ferroptosis in in vitro and in vivo models. Using a quantitative redox proteomics approach, we investigated the key regulated S-glutathionylated proteins in ferroptosis and found that the reduced S-glutathionylation of ADP-ribosylation factor 6 (ARF6) provided a labile iron pool to drive ferroptosis.

## RESULTS

### Reduced protein-SSG is associated with decreased glutathione pools by CHAC1 induction in multiple cell types undergoing ferroptosis

To investigate the pattern of protein-SSG in ferroptotic cells, we used the classical and commonly used ferroptosis inducer erastin because it can extensively induce ferroptosis in cells. In multiple human cell lines (human colonic adenocarcinoma [T84], hepatocellular carcinoma [HepG2], human oesophageal cancer [TE1], human prostate carcinoma [DU145], human lung adenocarcinoma [H1299], human ovarian cancer [A2780 and SKOV3], and human gastric cancer [HGC27]) undergoing ferroptosis induced by erastin (Supplementary Fig. 1a), the overall protein-SSG was significantly decreased after erastin treatment (Fig. 1a). Given that glutathione pools are strongly associated with protein-SSG, we measured the levels of GSH and GSSG in these cell lines and found that both were markedly depleted in the ferroptotic cells (Fig. 1b, Supplementary Fig. 1b). Considering that CHAC1 is responsible for glutathione degradation and GLRX is the most well-studied catalytic enzyme for deglutathionylation, we examined the expression levels of *CHAC1* and *GLRX* in these cells. *CHAC1* was significantly upregulated in all cell lines at both the mRNA and protein levels (Fig. 1c). We found that in some cells where protein-SSG was significantly decreased after erastin treatment, *GLRX* levels did not change in HepG2, TE-1, and H1299 cells. Even in cells, such as T84, DU145, HGC27, A2780, and SKOV3 with slightly higher *GLRX* levels, the induction of *CHAC1* was much more substantial than that of *GLRX* (Fig. 1c-d). These observations suggest that not mutually exclusive to GLRX-regulated deglutathionylation, CHAC1 regulated glutathione pool alterations may be key for protein-SSG in cell ferroptosis.

**Figure 1.**
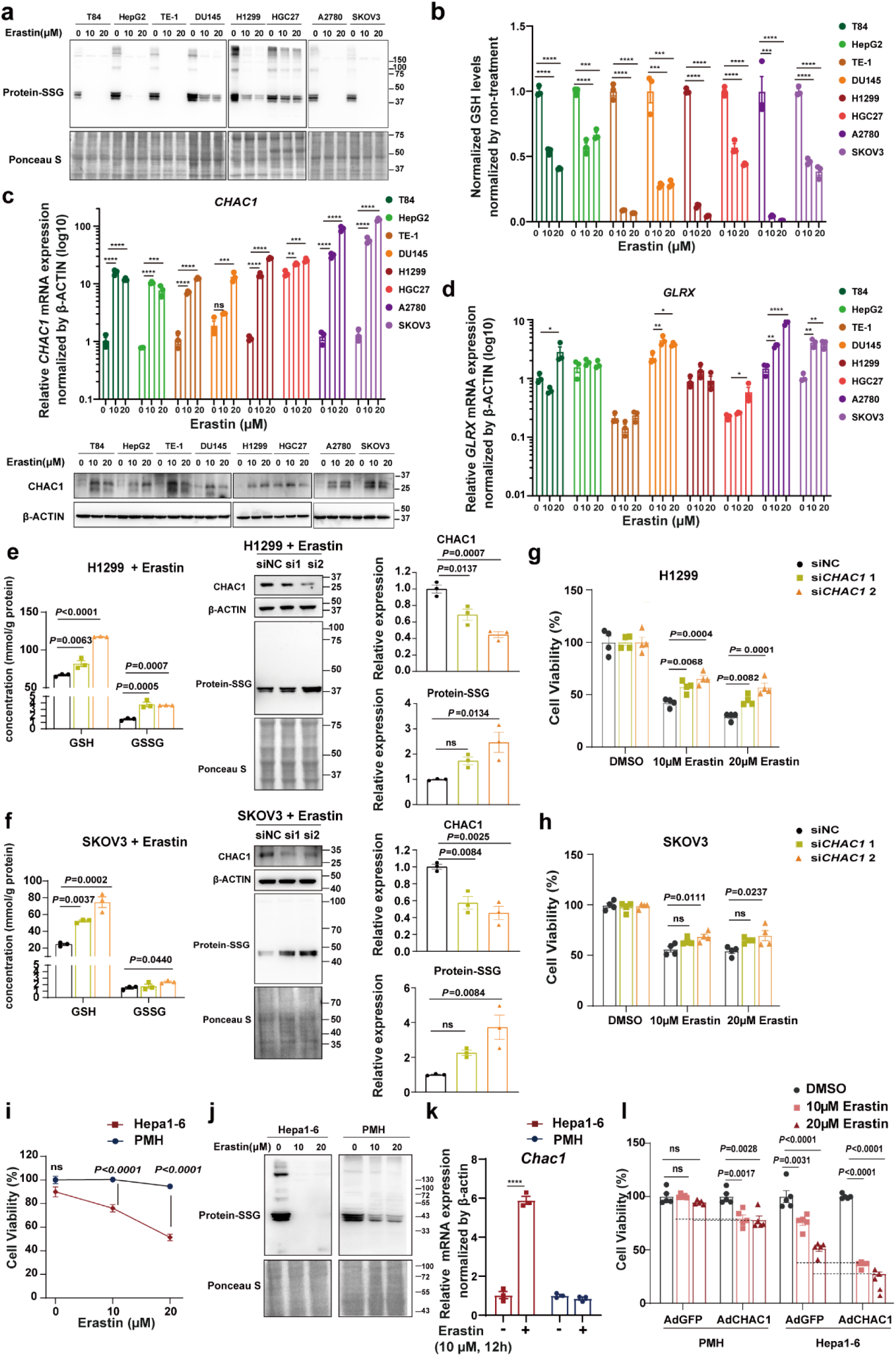
Reduced protein-SSG is associated with decreased glutathione pools by CHAC1 induction in multiple cell types undergoing ferroptosis. **a**: S-glutathionylated proteins in eight human cell lines treated with DMSO, or 10 μM or 20 μM erastin were analysed using western blotting with anti-glutathione. Ponceau S staining were used as internal references. **b**: Quantification of GSH and GSSG in eight human cell lines treated with DMSO, or 10 μM or 20 μM erastin. **c**: *CHAC1* mRNA and protein expression in eight human cell lines treated with DMSO or 10 μM, 20 μM erastin for 24 h were analysed using RT-qPCR and western blotting respectively. β-ACTIN were used as internal references for western blotting. **d**: *GLRX* mRNA expression in eight human cell lines treated with DMSO, or 10 μM or 20 μM erastin for 24 h. **e**: H1299 cells transfected with CHAC1 small interfering RNA (siRNA) and then treated with 10 μM erastin. The levels of GSH and GSSG were quantified by GSSG/GSH Quantification Kit. CHAC1 and S-glutathionylated proteins were analysed by Western blots and quantified. β-ACTIN and Ponceau S staining were used as internal references. **f**: SKOV3 cells transfected with CHAC1 siRNA and then treated with 10 μM erastin. The levels of GSH and GSSG were quantified by GSSG/GSH Quantification Kit. CHAC1 and S-glutathionylated proteins were analysed by Western blots and quantified. β-ACTIN and Ponceau S staining were used as internal references. **g**: Cell viability of H1299 cells transfected with *CHAC1* siRNA and then treated with DMSO, 10 μM, or 20 μM erastin for 24 h was measured by CellTiter-Glo® luminescent cell viability assay. **h**: Cell viability of SKOV3 cells transfected with *CHAC1* siRNA and then treated with DMSO, 10 μM, or 20 μM erastin for 24 h. **i**: Cell viability of PMHs and Hepa1-6 cells treated with DMSO, or 10 or 20 μM erastin for 24 h. **j**: S-glutathionylated proteins in PMHs and Hepa1-6 cells treated with DMSO or 10 μM, 20 μM erastin was analysed by Western blots. Ponceau S staining was used as an internal reference. **k**: *Chac1* mRNA expression in PMHs and Hepa1-6 cells treated with DMSO or 10 μM erastin for 12 h was analysed by RT-qPCR. **l**: Cell viability of Ad-GFP or Ad-CHAC1 PMH and Hepa1-6 treated with DMSO, or 10 μM or 20 μM erastin for 24 h. * *P* < 0.05, ** *P* < 0.01, *** *P* < 0.001, and **** *P* < 0.0001. PMH, primary mouse hepatocyte

We knocked down CHAC1 expression using siRNA in H1299 and SKOV3 cells exposed to erastin (Supplementary Fig. 1c) to confirm the importance of glutathione availability in protein S-glutathionylation. Unsurprisingly, the GSH and GSSG levels were increased by CHAC1 knockdown in ferroptotic H1299 and SKOV3 cells. Meanwhile, the overall SSG protein levels were increased by *CHAC1* siRNA in elastin-treated H1299 (Fig. 1e) and SKOV3 cells (Fig. 1f). To explore the effect of decreased glutathione pools on ferroptosis; we evaluated the viability of H1299 and SKOV3 cells after elastin treatment in the presence or absence of CHAC1 siRNA transfection. As a result, the knockdown of CHAC1 restrained cell death in both H1299 (Fig. 1g) and SKOV3 cells exposed to elastin (Fig. 1h).

We also tested this phenomenon in such primary cells as primary mouse hepatocytes (PMHs). Surprisingly, PMHs were not responsive to erastin. In comparison, Hepa1-6 cells (mouse hepatocellular carcinoma cells) were sensitive to erastin (Fig. 1i). We were curious whether this discrepancy was related to the differences in protein-SSGs. Erastin inhibited almost all protein-SSG in Hepa1-6 cells (Fig. 1j); however, protein-SSG was still observed in PMHs. Accordingly, upon erastin treatment, CHAC1 was not induced in PMHs but was substantially upregulated in Hepa1-6 cells (Fig. 1h). Overexpression of CHAC1 resulted in decreased cell viability when challenged with erastin compared to the control in both PMHs and Hepa1-6 cells (Fig. 1l).

Together, using multiple cell types and erastin-induced ferroptosis, we found that CHAC1-related glutathione pools and protein S-glutathionylation are broadly involved in cell ferroptosis.

### Decreased glutathione pools by CHAC1 reduce protein-SSG and aggravate APAP-induced hepatotoxicity and ferroptosis in APAP-injured mice liver

We tried to introduce an in vivo model to investigate the role of the CHAC1-mediated glutathione pool and protein-SSG reduction in ferroptosis. Our lab and other groups revealed that hepatocyte ferroptosis substantially contributes to APAP-induced acute liver injury in mouse models (Lorincz, Jemnitz et al., 2015, Niu, Lei et al., 2022, Yamada, Karasawa et al., 2020), so we revisited our published RNA sequencing data ((Li, Ming et al., 2018), SRA database of NCBI: PRJNA731100) and found that *Chac1* is one of the top upregulated genes in liver tissues from mice challenged with 300 or 750 mg/kg APAP (Fig. 2a, Supplementary Fig. 2a, b). The significant induction of *Chac1* mRNA in the liver tissues of mice 3, 6, and 12 h after 300 mg/kg APAP challenge was validated using RT-qPCR (Fig. 2b). The mRNA level of *Chac1* was also dose-dependently induced by APAP challenge in PMHs (Fig. 2c). We detected CHAC1 expression in the liver tissues of patients with APAP-induced liver injury (AILI) and healthy controls using IHC. Consistently, CHAC1 expression (presented as H-scores in IHC sections) was significantly higher in liver tissues from patients with AILI than in those from healthy controls (Fig. 2d).

**Figure 2.**
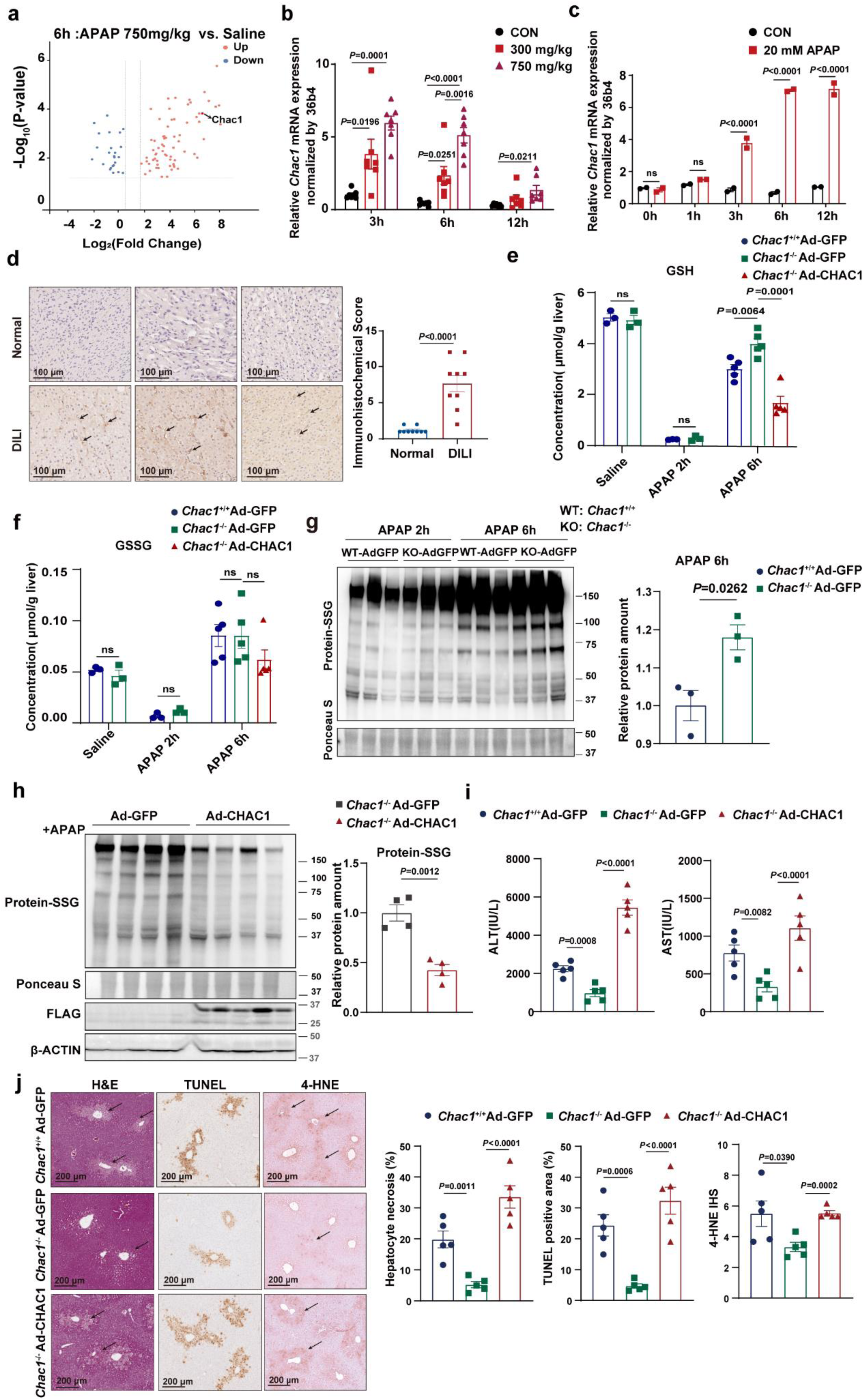
Decreased glutathione pools by CHAC1 reduce protein-SSG and aggravate APAP-induced hepatotoxicity and ferroptosis in APAP-injured mice liver. **a**: Volcano plots shows upregulated and downregulated genes from RNA transcriptome data of group treated with 750 mg/kg APAP for 6 h compared to the saline group (Fold change ≥ 1.5, *P* < 0.05; n = 3 mice/group). **b**: *Chac1* mRNA expression in mouse liver tissues treated with saline or 300 and 750 mg/kg APAP for 3, 6, or 12 h. **c**: *Chac1* mRNA expression in PMHs with or without 20 mM APAP challenge for 1, 3, 6, and 12 h was analysed by RT-qPCR. **d**: Immunohistochemical staining of CHAC1 in liver sections from healthy controls (*n* = 9) and patients with AILI (*n* = 9), followed by immunohistochemical score of liver tissue. The black arrow indicates positive staining. Scale bar = 100 μm. **e, f**: Quantification of GSH and GSSG in the liver tissues of *Chac1^+^*^/+^ Ad-GFP, *Chac1^-^*^/-^ Ad-GFP, and *Chac1^-^*^/-^ Ad-CHAC1 mice treated with saline or 300 mg/kg APAP for 2 and 6 h. **g**: S-glutathionylated proteins in the liver tissues of *Chac1^+^*^/+^ Ad-GFP and *Chac1^-^*^/-^ Ad-GFP mice treated with 300 mg/kg APAP for 2 and 6 h were analysed by western blots with anti-glutathione. Ponceau S staining was used as an internal reference. The relative expression levels of S-glutathionylated proteins after APAP treatment for 6 h were quantified. **h**: S-glutathionylated proteins and CHAC1-FLAG protein in the liver tissues of *Chac1^-^*^/-^ Ad-GFP and *Chac1^-^*^/-^ Ad-CHAC1 mice treated with 300 mg/kg APAP for 6 h were analysed by western blotting. **i**: Serum levels of ALT and AST in *Chac1^+^*^/+^ Ad-GFP, *Chac1^-^*^/-^ Ad-GFP, and *Chac1^-^*^/-^ Ad-CHAC1 mice treated with saline or 300 mg/kg APAP for 6 h. **j**: H&E staining, TUNEL staining and 4-HNE protein adducts staining in liver tissues of *Chac1^+^*^/+^ Ad-GFP, *Chac1^-^*^/-^ Ad-GFP and *Chac1^-^*^/-^ Ad-CHAC1 mice treated with 300 mg/kg APAP for 6 h. Scale bars = 200 μm. The hepatocyte necrosis area, TUNEL-positive area, and immunohistochemical score for 4-HNE protein adduct staining were quantified. AILI, APAP-induced liver injury; ALT, alanine aminotransferase; APAP, acetaminophen; AST, aspartate aminotransferase; H&E, haematoxylin and eosin; PMH, primary mouse hepatocyte

We generated *Chac1*^+/+^ and *Chac1*^-/-^ mice (Supplementary Fig. 2c, d) to investigate the effect of upregulated CHAC1 on glutathione pools, protein-SSG, and ferroptosis during AILI, and found that they exhibited normal growth and development. The mRNA level of *Chac1* in the liver was time-dependently induced by APAP in *Chac1*^+/+^ mice, but not in *Chac1*^-/-^ mice (Supplementary Fig. 2e-f). We also rescued CHAC1 expression in *Chac1*^-/-^ mice by injecting Ad-CHAC1 via the tail vein. The mice were challenged with APAP. As a result, we found that the levels of GSH and GSSG did not differ between *Chac1*^-/-^and *Chac1*^+/+^ mice in terms of homeostatic status. Upon APAP administration, the concentrations of both GSH and GSSG in the liver markedly decreased after 2 h and then started to increase in mice of both genotypes (Figs. 2e and f and Supplementary Fig. 3a). Notably, the level of GSH was restored faster in *Chac1*^-/-^ mice, than in *Chac1*^+/+^ mice, after 6 h (Fig. 2e). As predicted, although the overall protein-SSG in the liver tissues of mice of both genotypes decreased after 2 h, it increased 6 h after APAP administration (Fig. 2g, Supplementary Fig 3b). At 6 h, the levels of protein-SSG in the livers of *Chac1*^-/-^ mice were higher than those in *Chac1*^+/+^ mice (Fig. 2g), suggesting that CHAC1 deficiency accelerated the recovery of liver glutathione depletion and increased liver protein-SSG levels in AILI mice. Moreover, the concentrations of glutathione and the levels of protein-SSG in the liver were decreased by the overexpression of CHAC1 in *Chac1*^-/-^ mice (Figs. 2e, f, and h, and Supplementary Fig. 3a), suggesting that CHAC1 overexpression retarded the recovery of liver glutathione depletion and reduced liver protein-SSG in AILI mice.

Next, we assessed the effects of the CHAC1-mediated glutathione pool and protein-SSG reduction on APAP-induced hepatotoxicity and ferroptosis. As a result, the Serum levels of ALT and AST were significantly decreased in *Chac1*^-/-^ mice than those in *Chac1*^+/+^ mice following the administration of 300 mg/kg APAP for 6 h (Fig. 2i). Hematoxylin and eosin (H&E) and Terminal dUTP nick-end labeling (TUNEL) staining revealed that *Chac1* deficiency significantly decreased the APAP-induced centrilobular hepatic necrosis and DNA fragmentation (Fig. 2j). We tested whether CHAC1 contributed to hepatocyte ferroptosis. Staining with 4-hydroxynonenal (4-HNE), a reliable indicator of endogenous lipid peroxidation in vivo, revealed that compared with *Chac1*^+/+^ mice, *Chac1*^-/-^ mice showed a clear reduction in 4-HNE-positive cells in liver histological sections (Fig. 2j). Further, we found that the improvement of hepatotoxicity in *Chac1*^-/-^ mice could be retarded by CHAC1 overexpression (Fig. 2i, j).

The decreased glutathione pools induced by CHAC1 reduced protein-SSG levels and aggravated APAP-induced hepatotoxicity and ferroptosis in APAP-injured mouse livers.

### Decreased glutathione pools by CHAC1 reduce protein-SSG and aggravate APAP-induced hepatotoxicity and ferroptosis in primary mouse hepatocytes

We isolated and cultured PMHs from *Chac1*^+/+^ and *Chac1*^-/-^ mice to further verify the strong correlation among glutathione pools, protein-SSG, and ferroptosis at the cellular level. Deletion of CHAC1 in PMHs increased SSG protein levels (Fig. 3a). Next, we studied the effects of CHAC1 on glutathione pools and protein-SSGs in wild-type PMHs using Ad-CHAC1 or Ad-GFP adenoviruses at different multiplicity of infection values (0.1, 1, 2.5, 5, and 10). After 12 or 36 h of infection (Fig. 3b, c, Supplementary Fig. 3c, d, e), the PMHs were treated with or without 20 mM APAP for 6 h. As a result, CHAC1 overexpression dose-dependently reduced protein-SSG in APAP-challenged PMHs (Fig. 3b, Supplementary Fig. 3c). Meanwhile, in the Ad-GFP group, upon APAP challenge, GSH was depleted, the GSSG content increased, and the GSH/GSSG ratio decreased. However, changes in GSH and GSSG levels upon APAP challenge were significantly diminished in the Ad-CHAC1 group, possibly because both GSH and GSSG levels were extremely low after CHAC1 overexpression. CHAC1 overexpression dramatically decreased the GSH and GSSG content in PMHs in a dose-dependent manner (Fig. 3c, Supplementary Fig. 3d, e).

**Figure 3.**
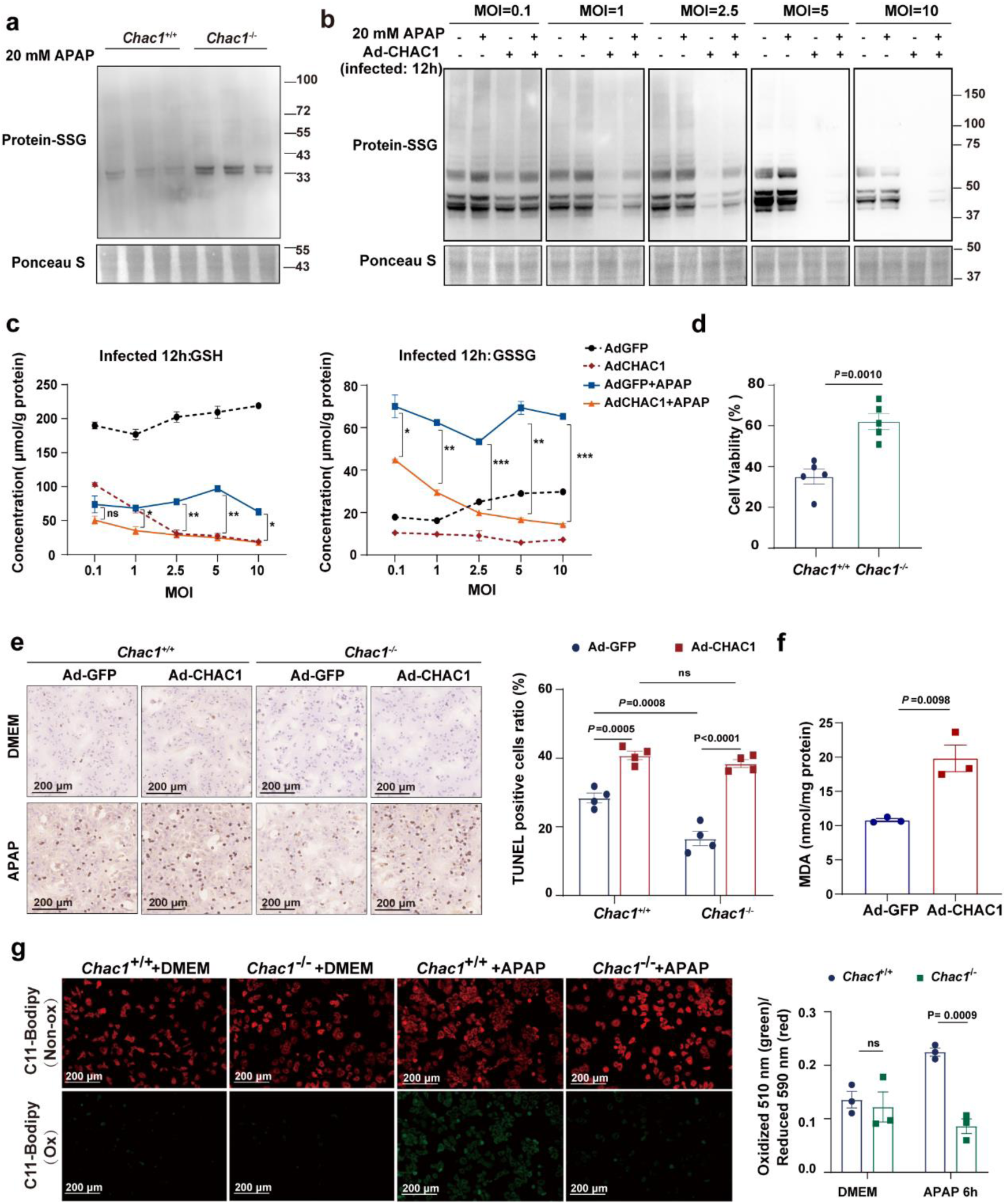
Decreased glutathione pools by CHAC1 reduce protein-SSG and aggravate APAP-induced hepatotoxicity and ferroptosis in primary mouse hepatocytes. **a**: S-glutathionylated proteins in PMHs from *Chac1^+^*^/+^ and *Chac1^-^*^/-^ mice treated with 20 mM APAP for 3 h were analysed by western blotting with anti-glutathione. Ponceau S staining was used as an internal reference. **b**: S-glutathionylated proteins in PMHs infected with Ad-GFP or Ad-CHAC1 at a multiplicity of infection (MOI)=0.1, 2.5, 5, and 10 for 12 h and then treated with 20 mM APAP for 6 h were analysed by western blotting with anti-glutathione. Ponceau S staining was used as an internal reference. **c**: Quantification of GSH and GSSG in PMHs infected with Ad-GFP or Ad-CHAC1 adenovirus at MOI=0.1, 2.5, 5, and 10 for 12 h and then treated with 20 mM APAP for 6 h (*n* = 2, * *P* < 0.05, ** *P* < 0.01, *** *P* < 0.001, and **** *P* < 0.0001 via *t* test). **d:** PMHs were isolated from *Chac1^+^*^/+^ and *Chac1^-^*^/-^ mice and treated with 20 mM APAP for 12 h. Cell viability was measured using the CellTiter-Glo luminescent cell viability assay (*n* = 5, *t* test). **e**: Isolated *Chac1^+^*^/+^ and *Chac1^-^*^/-^ PMHs were infected at MOI of 10 for 36 h and then treated with 20 mM APAP. After 6 h, TUNEL staining was performed for nuclear DNA fragmentation (scale bars = 200 μm). **f**: Determination of malondialdehyde (MDA) levels in PMHs infected with Ad-GFP or Ad-CHAC1 adenoviruses at MOI=10 for 36 h and then treated with 20 mM APAP for 12 h. **g**: PMHs from *Chac1^+^*^/+^ and *Chac1^-^*^/-^ mice were treated with 20 mM APAP for 12 h. Representative images of C11 Bodipy 581/591 fluorescence. Red fluorescence represents non-lipid oxidation, and green fluorescence represents lipid oxidation. The statistical chart shows the ratio of green to red fluorescence (scale bars = 200 μm). APAP, acetaminophen; GSH, glutathione; GSSG, oxidised glutathione; PMH, primary mouse hepatocyte

Next, we assessed the effects of the CHAC1-mediated glutathione pool and protein-SSG reduction on APAP-induced hepatotoxicity and ferroptosis. The viability of PMHs was significantly increased upon APAP challenge in the *Chac1*^-/-^ group compared with that in the *Chac1*^+/+^ group (Fig. 3d). TUNEL staining revealed that CHAC1 deficiency was attenuated whereas the overexpression aggravated APAP-induced hepatotoxicity (Fig. 3e). Upon APAP challenge, overexpression of CHAC1 increased the levels of the lipid peroxidation product malondialdehyde (MDA) in *Chac1*^-/-^ PMHs (Fig. 3f). BODIPY 581/591 C11 (BODIPY) probes are commonly used to detect lipid peroxidation in living cells. Upon oxidation, the fluorescence of the probe shifts from red to green. Consistently, lipid peroxidation was markedly decreased in *Chac1*^-/-^ PMHs compared with that in *Chac1*^+/+^ PMHs (Fig. 3g).

The association between SSG and ferroptosis was clearly observed in vivo and in vitro. The decreased glutathione pools caused by CHAC1 reduced protein-SSG levels and aggravated APAP-induced hepatotoxicity and ferroptosis in primary mouse hepatocytes.

### Mass-spectroscopic quantification of CHAC1-regulated protein cysteine S-glutathionylation in ferroptotic PMHs induced by APAP

We confirmed that cysteine S-glutathionylation is regulated by CHAC1-induced glutathione pool depletion in multiple cell lines and hepatocytes. Therefore, we conducted a quantitative redox proteomic analysis to identify the S-glutathionylated proteins involved in APAP-induced hepatocyte ferroptosis in the presence and absence of CHAC1 overexpression. PMHs for glutathionylomic analysis were infected with Ad-GFP or Ad-CHAC1 for 36 h and then challenged with 20 mM APAP for 3 h (Fig. 4a). We identified 1105 unique glutathionylated peptides from 482 proteins (Fig. 4b). Oxidative state induced by APAP challenge markedly enhanced (Fold change ≥ 1.2, *P* < 0.05) cysteine S-glutathionylation in both Ad-GFP and Ad-CHAC1 groups (Fig. 4c, d). Compared with the Ad-GFP group, CHAC1 overexpression markedly decreased cysteine S-glutathionylation, especially after APAP treatment, which was consistent with the western blot data (Fig. 4c, e). APAP induced more S-glutathionylation at 221 cysteine sites, which was decreased by CHAC1 overexpression (Fig. 4f, g). GO analysis revealed that these proteins were enriched in biological processes, such as lipid metabolism, mitochondrial respiratory chain, and arachidonic acid metabolism, which are all potentially relevant to hepatotoxicity and ferroptosis (Supplementary Fig. 4).

**Figure 4.**
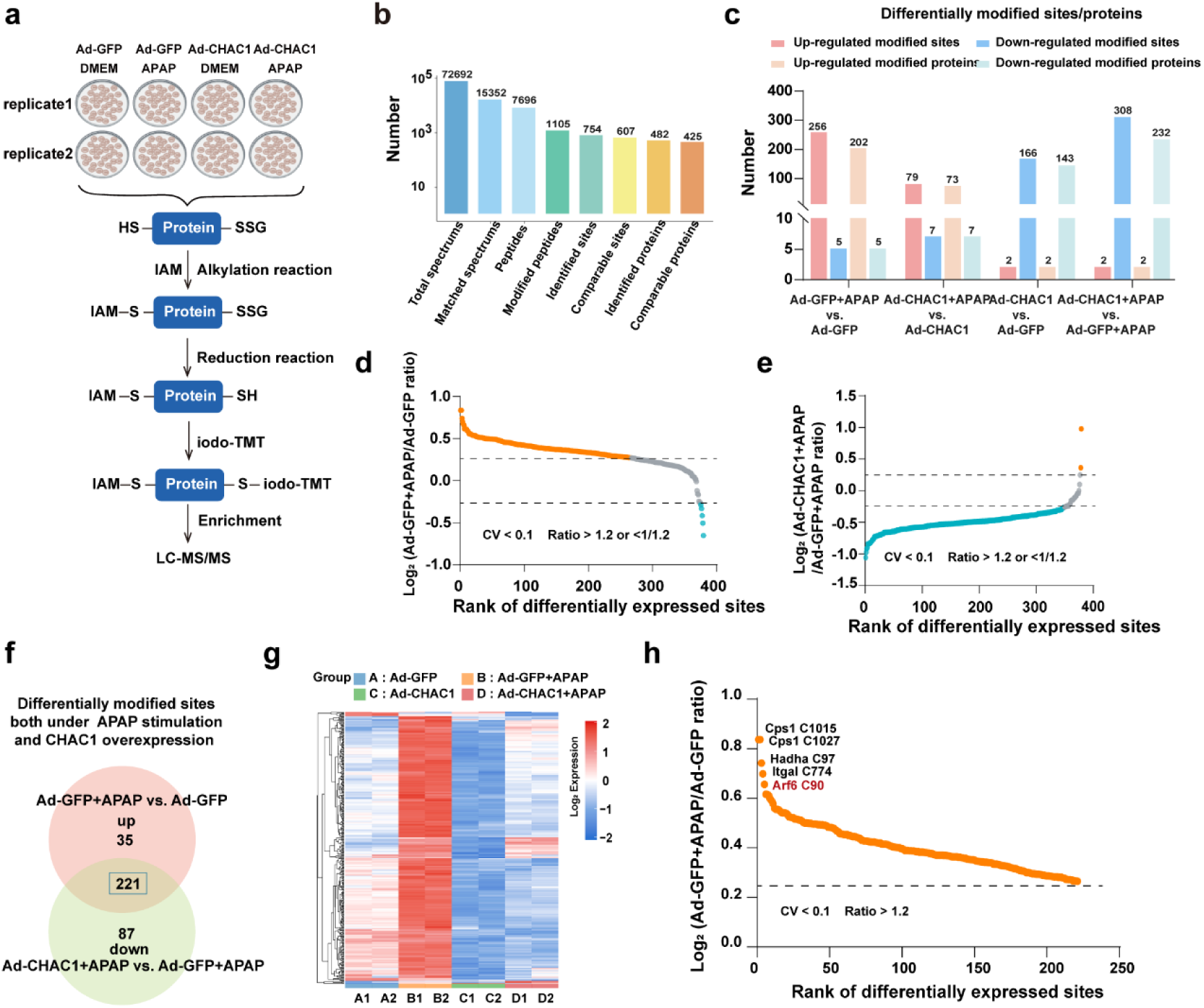
Mass-spectroscopic quantification of CHAC1-regulated protein cysteine S-glutathionylation in ferroptotic PMHs induced by APAP. **a**: Flowchart outlining the key experimental procedures for proteomic analysis of S-glutathionylation. **b**: Overview of the identification of modification sites. **c**: The histogram shows the distribution of differential modification sites/proteins among different comparison groups (Coefficient of Variation (CV) < 0.1, fold change ≥ 1.2). **d**: Scatter plot showing the distribution of differential modification sites sorted by the ratio of Ad-GFP + APAP/Ad-GFP. Red dots indicating upregulation of significant differences, blue dots indicating significant differences down-regulation, and grey indicating no significant differences (CV < 0.1, fold change ≥ 1.2). **e**: Scatter plot showing the distribution of differential modification sites sorted by the ratio of Ad-CHAC1 + APAP/Ad-GFP + APAP. Red dots indicating up-regulation of significant differences, blue dots indicating significant differences down-regulation, and grey indicating no significant differences (CV < 0.1, fold change ≥ 1.2). **f**: Venn diagram shows differentially modified sites both under APAP stimulation and CHAC1 overexpression (Fold change ≥ 1.2). **g**: The heatmap shows the union of differential modification sites in Ad-GFP, Ad-GFP + APAP, Ad-CHAC1, and Ad-CHAC1 + APAP comparison groups (CV < 0.1, fold change ≥ 1.2). **h**: The scatter plot shows differentially modified sites both under APAP stimulation and CHAC1 overexpression; the order was sorted by the ratio of Ad-GFP + APAP / Ad-GFP (CV < 0.1, fold change ≥ 1.2). PMH, primary mouse hepatocyte

### Cys90 S-glutathionylation of ARF6 promotes ARF6 activation and alleviates ferroptosis

Among the proteins with decreased S-glutathionylation due to CHAC1 overexpression, we found that ARF6 is one of the top five proteins upregulated by APAP treatment in PMHs (Fig. 4h). ARF6 is a member of the small GTPase ADP-ribosylation factor (ARF) family that converts inactive GDP-binding to active GTP-binding conformations. According to the redox proteomic analysis, ARF6 is glutathionylated on (Cys90) (Fig. 5a). We found that glutathionylation in Cys90 of the ARF6 protein increased upon APAP challenge in Ad-GFP PMHs, but not in Ad-CHAC1 PMHs (Fig 5b).

**Figure 5.**
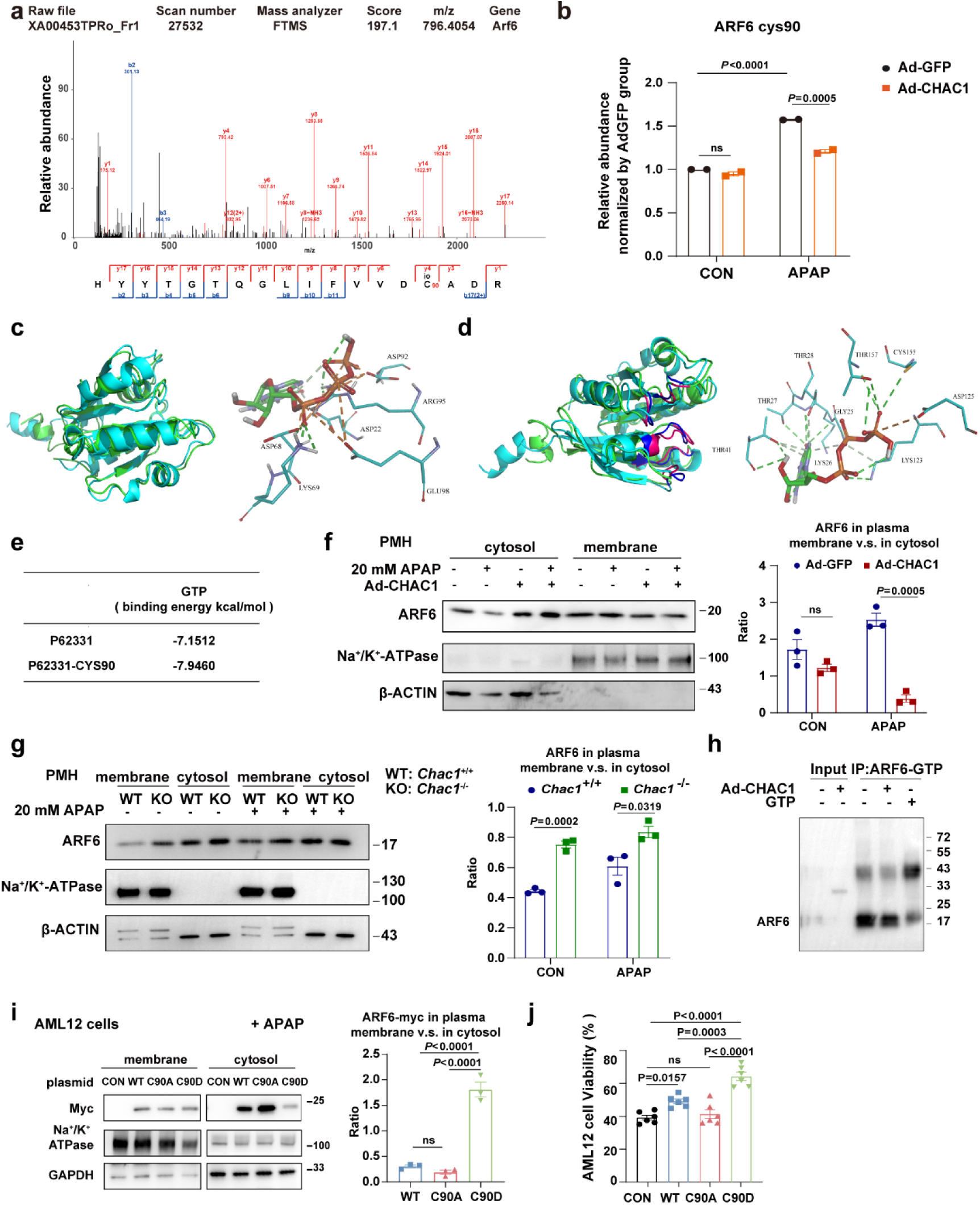
Cys90 S-glutathionylation of ARF6 promotes ARF6 activation and alleviates ferroptosis. **a**: Two-stage mass spectrometry of the glutathionylated peptide from ARF6. The secondary mass spectrum shows fragment ion information of the ARF6 C90 peptide segment. **b**: Histogram showing the relative modification abundance of ARF6 C90 in different treatment groups, with glutathionylated peptides identified and quantified by LC-MS/MS (all values were standardised by the mean of the AdGFP-CON group). **c**: Illustrations of predicted 3D structures of ARF6 and Cys90 glutathionylated ARF6 from PyMOL. Structural alignment of modelled ARF6 and modelled Cys90 glutathionylated ARF6 (cyan: before modification; green: after modification). **d**: Regions with significant changes in the predicted 3D structures of ARF6 and Cys90 glutathionylated ARF6 from PyMOL. Structural alignment of the modelled ARF6 and Cys90 glutathionylated ARF6 (blue: before modification; red: after modification). **e**: Docking score between ARF6 and GTP and the docking score between Cys90 glutathionylated ARF6 and GTP. **f**: PMHs were isolated from C57BL/6 mice and infected with Ad-CHAC1 or Ad-GFP at an MOI of 10 for 36 h, followed by treatment with 20 mM APAP for 6 h. Proteins were extracted and fractionated into cytosolic and membrane fractions. The ARF6 protein was analysed using western blotting. β-ACTIN marks the cytosol, and Na^+^/K^+^-ATPase marks the membrane. Statistical chart shows the ratio of ARF6 in the plasma membrane to ARF6 in the cytosol. **g**: PMHs were isolated from *Chac1^+^*^/+^ or *Chac1^-^*^/-^ mice and treated with 20 mM APAP for 6 h. Proteins were extracted and fractionated into cytosolic and membrane fractions. Protein ARF6 was analysed using western blotting. β-ACTIN marks the cytosol, and Na^+^/K^+^-ATPase marks the membrane. Statistical chart shows the ratio of ARF6 in the plasma membrane to ARF6 in the cytosol. **h**: Pull-down assay showing the expression of activated ARF6 in Ad-CHAC1 or Ad-GFP PMHs treated with 20 mM APAP for 6 h (pull-down, ARF6-GTP; IB, ARF6). Whole cell lysates to confirm the expression of ARF6. **i**: Expression of the ARF6-Myc-tag protein in AML12 cells transfected with CON, WT, C90A, or C90D plasmids and treated with APAP for 6 h was analysed using western blotting. GAPDH marks the cytosol and Na^+^/K^+^-ATPase marks the membrane. Statistical chart shows the ratio of ARF6 in the plasma membrane to ARF6 in the cytosol. **j**: The viability of AML12 cells transfected with CON, WT, C90A, or C90D plasmids and treated with APAP for 12 h was quantified using the CellTiter-Glo luminescent cell viability assay. PMH, primary mouse hepatocyte

ARF6 has a highly conserved sequence across species. The protein sequence of ARF6 is identical in mice and humans. We used a molecular modelling approach to determine whether glutathionylation at Cys90 affected the binding energy between ARF6 and GTP. The structure of the ARF6 protein was predicted using AlphaFold 2, which was used for subsequent protein docking and structural modifications. The reliability of the ARF6 model was validated by analysing its Ramachandran plot (Supplementary Fig. 5a). A total of 23 binding sites on the ARF6 model protein were predicted using the MOE 2015 software. The reliability of glutathionylated ARF6 in the Cys90 model was validated by analysing its Ramachandran plot (Supplementary Fig. 5b). The structures of ARF6 and Cys90 glutathionylated ARF6 were viewed in PyMOL, which showed substantial changes in the structures (RMSD=1.312) after Cys90 glutathionylation compared to the original ARF6 (cyan) (Fig. 5c). Regions with greater changes were further analysed. The glutathionylation of Cys90 had a significant impact on the spatial position of the active site (Fig. 5d). GTP/GDP had a binding free energy of -7.1512/-6.7329 kcal/mol for ARF6 and a binding free energy of -7.9460/-7.8688 kcal/mol for Cys90 glutathionylated ARF6 (Fig. 5e). Docking data suggested that glutathionylation at Cys90 promoted the binding of GTP/GDP to ARF6, which may have increased ARF6 activity.

ARF6 cycles between the plasma membrane (activated form, ARF6-GTP) and the endosomal compartment (inactivated form, ARF6-GTP) (Radhakrishna & Donaldson, 1997). We separated the membrane and cytoplasmic proteins from Ad-GFP- or Ad-CHAC1-infected PMHs in the presence or absence of APAP. We found that APAP increased, but overexpression of CHAC1 decreased, membrane-localised ARF6 (Fig. 5f). We then separated the membrane and cytoplasmic proteins from *Chac1*^+/+^ and *Chac1*^-/-^ PMHs with or without APAP challenge. Consistently, APAP increased, whereas CHAC1 deficiency enhanced membrane localisation of ARF6 (Fig. 5g). Next, we performed a pull-down assay with an antibody specific to ARF-GTP-conjugated agarose and examined its levels by western blot analysis. CHAC1 promotes ARF6 GTP loading (Fig. 5h). These findings suggest that CHAC1 decreased ARF6 membrane localization and inactivated the membrane localisation of ARF6.

To examine whether Cys90 glutathionylation is involved in AFR6 activation, we constructed AML-12 cells in which ARF6-WT and its mutants (ARF6-C90A and ARF6-C90D) were overexpressed with C90A to mimic the unglutathionylated form of ARF6 and the C90D mutant to mimic the glutathionylated form of ARF6. As expected, the examination of ARF6 localisation using western blotting revealed that the ratio of ARF6 in the plasma membrane to ARF6 in the cytosol decreased when ARF6-C90A-myc was overexpressed, whereas it significantly increased when ARF6-C90D-myc was overexpressed. This suggests that glutathionylation increases ARF6 membrane localization and ARF6 activation (Fig. 5i). Furthermore, overexpression of ARF6-C90A aggravated, but overexpression of ARF6-C90D suppressed, APAP-induced hepatotoxicity in AML12 cells (Fig. 5j).

These results indicate that glutathione degradation catalysed by CHAC1 decreased ARF6 S-glutathionylation at Cys90. S-glutathionylation of ARF6 promotes its activation and alleviates ferroptosis in hepatocytes.

### CHAC1 delayed endosomal recycling of transferrin receptors and increased the labile iron pool to promote APAP-induced ferroptosis in PMHs

A few reports mentioned that ARF6 suppressed ferroptosis in pancreatic and gastric cancer cells (Geng & Wu, 2022, Ye, Hu et al., 2020). To determine whether ARF6 plays a role in APAP-induced hepatocyte ferroptosis, we knocked down ARF6 in PMHs using siRNA and treated the cells with APAP (Fig. 6a). We found that both CellTiter-Glo® Luminescent Cell Viability assays and Live & Dead cell stanning assays revealed the aggravated cell death following the ARF6 knock down in PMHs exposed to APAP overdose (Fig. 6b, Supplementary Fig. 6a). Lipid peroxidation was markedly increased by *ARF6* siRNA compared to that with control siRNA (Fig. 6c, Supplementary Fig. 6b). We then detected the labile iron pool using the FerroOrange software. ARF6 knockdown significantly increased intracellular Fe^2+^ levels in PMHs upon APAP challenge (Fig. 6d, Supplementary Fig. 6c). These results suggested a protective role for ARF6 in APAP-induced hepatocyte ferroptosis.

**Figure 6.**
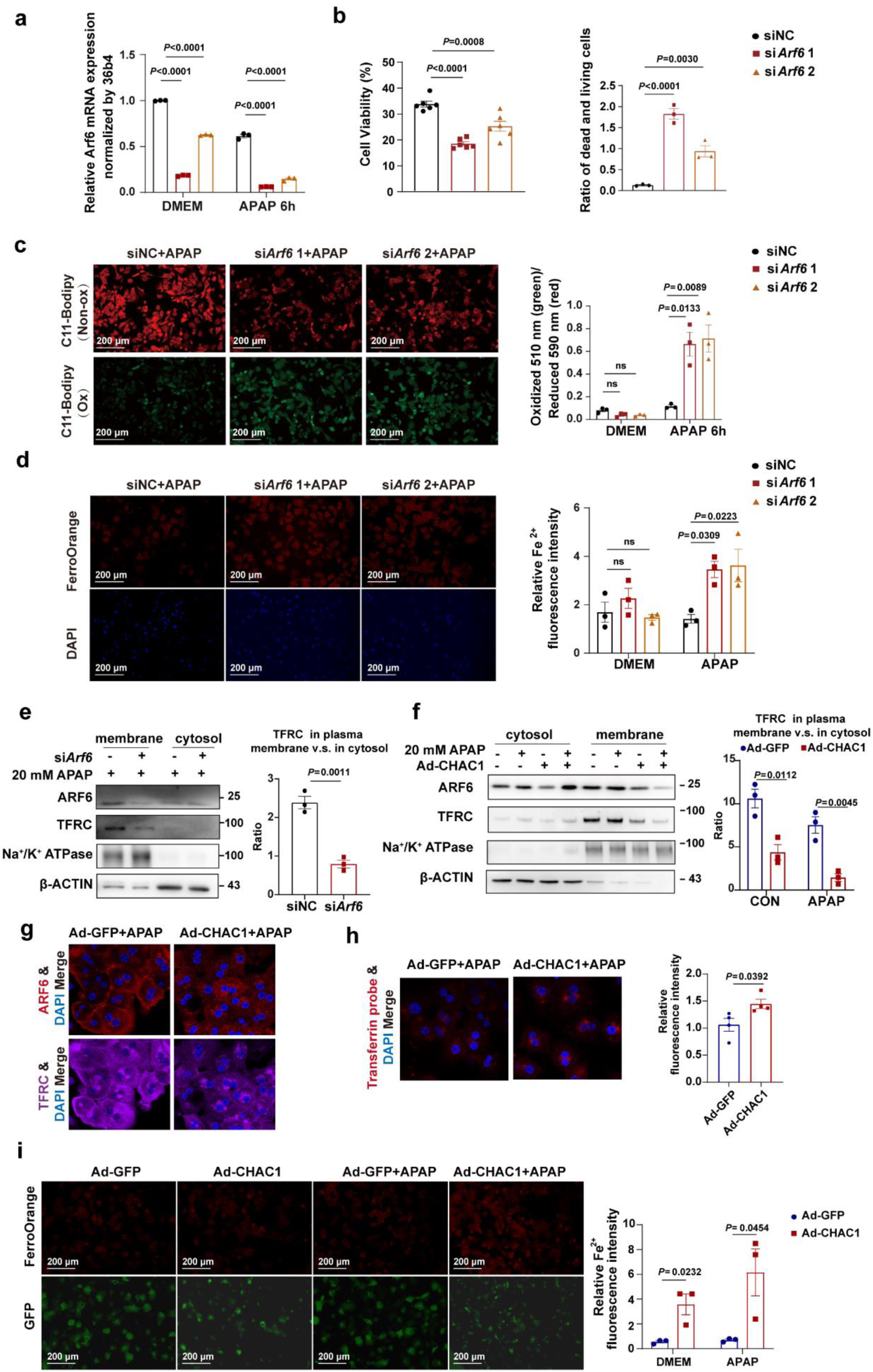
CHAC1 delayed endosomal recycling of transferrin receptors and increased the labile iron pool to promote APAP-induced ferroptosis in PMHs. **a**: *Arf6* mRNA expression in PMHs transfected with *Arf6* siRNA for 48 h and then treated with DMEM or 20 mM APAP, as analysed by RT-qPCR. **b**: PMHs were transfected with *Arf6* siRNA and treated with 20 mM APAP. Cell viability was measured using the CellTiter-Glo luminescent cell viability assay. The ratio of dead to live cells was measured using calcein-AM/PI live/dead cell staining. **c**: Representative images of C11 Bodipy 581/591 fluorescent probe, which was used for detecting the formation of lipid peroxides (Red: non-lipid oxidation; green: lipid oxidation, Scale bars = 200 μm). The statistical chart shows the ratio of green to red fluorescence. **d**: Representative images of the FerroOrange fluorescent probe used to detect labile ferrous ions (Red: FerroOrange; blue: DAPI, Scale bars = 200 μm). The statistical chart shows the relative Fe^2+^ fluorescence intensity. **e**: ARF6 and TFRC protein expression in PMHs transfected with *Arf6* siRNA and treated with 20 mM APAP was analysed using western blotting. β-ACTIN marks the cytosol, and Na^+^/K^+^-ATPase marks the membrane. The statistical chart shows the ratio of TFRC in the plasma membrane to TFRC in the cytosol. **f**: ARF6 and TFRC protein expression in PMHs infected with Ad-GFP or Ad-CHAC1 adenovirus and treated with 20 mM APAP was analysed by western blotting. β-ACTIN marks the cytosol, and Na^+^/K^+^-ATPase marks the membrane. The statistical chart shows the ratio of TFRC in the plasma membrane to TFRC in the cytosol. **g**: Immunofluorescence staining (with ARF6 and TFRC) in PMHs infected with Ad-GFP or Ad-CHAC1 adenovirus and then treated with 20 mM APAP (Red: ARF6; purple: TFRC; blue: DAPI, Scale bars = 200 μm). **h**: Representative images of the transferrin fluorescent probe (red: transferrin; blue: DAPI, Scale bars = 200 μm). The statistical chart shows the relative transferrin content. **i**: Representative images of the FerroOrange fluorescent probe (Red: FerroOrange; green: GFP, Scale bars = 200 μm). The statistical chart shows the relative Fe^2+^ fluorescence intensity. PMH, primary mouse hepatocyte

ARF6 regulates transferrin receptor (TFRC) recycling in Chinese hamster ovary cells (D’Souza-Schorey, Li et al., 1995). We speculated that ARF6 participates in TFRC recycling during ferroptosis. Western blot analysis revealed that ARF6 knockdown decreased TFRC protein levels in the plasma membrane (Fig. 6e). Next, we investigated the effects of CHAC1 on endosomal recycling of TFRC. In the presence of APAP, the protein level of TFRC in the plasma membrane decreased, which is in line with ARF6 membrane localization (Fig. 6f). To further confirm this, we performed immunofluorescence staining for ARF6 and TFRC in PMHs. Upon APAP challenge, overexpression of CHAC1 resulted in a decrease in cell surface binding of ARF6 and an increase in the endosomal localisation of ARF6 and TFRC as compared to that in control cells infected with the control-GFP virus (Fig. 6g). We found that CHAC1 overexpression increased the intracellular Fe^2+^ levels in PMHs upon APAP challenge (Fig. 6h).

Our data suggest that CHAC1, responsible for catalysing glutathione degradation and reducing ARF6 S-glutathionylation at Cys90, delays endosomal recycling of transferrin receptors and increases the labile iron pool to promote APAP-induced ferroptosis by inactivating ARF6 in PMHs.

### ARF6 glutathionylation at Cys90 promoted ARF6 activation and contributed to erastin-induced ferroptosis in multiple cell types

Regarding the dramatic change of protein glutathionylation widely occurred in multiple cell types undergoing ferroptosis as shown in Fig. 1, we transfected ARF6-WT and its mutant (ARF6-C90A and ARF6-C90D) in 293T and H1299 cells. C90A mimics the unglutathionylated form of ARF6, whereas the C90D mutant mimics the glutathionylated form of ARF6. In line with the results obtained in AML12 cells (Fig. 5j) undergoing APAP-induced ferroptosis, the ratio of ARF6 in the plasma membrane to ARF6 in the cytosol decreased when ARF6-C90A-myc was overexpressed, whereas it significantly increased when ARF6-C90D-myc was overexpressed in both 293T (Fig. 7a) and H1299 cells (Fig. 7b). Moreover, overexpression of ARF6-C90A promoted ferroptosis, but overexpression of ARF6-C90D alleviated ferroptosis induced by erastin in H1299 cells (Fig. 7c). To further observe the molecular alterations caused by ARF6 glutathionylation at Cys90, we performed RNA-seq using ARF6-myc (WT) or ARF6-C90A-myc transfected H1299 cells, with or without erastin treatment. Using GO or KEGG pathway enrichment analyses, we found that the 443 genes upregulated only in the ARF6-C90A group but not in the WT group were enriched in cholesterol biosynthesis, lipid metabolic process, autophagy, and ferroptosis. However, the 853 genes downregulated only in the ARF6-C90A group but not in the WT group were mainly involved in RNA splicing, cell division, cell cycle, DNA repair, and endocytosis. The suppression of endocytosis caused by ARF6 glutathionylation at Cys90 was also reflected by GO analyses on 720 genes upregulated only in WT group but not in the ARF6-C90A group (Fig. 7d). These findings suggest that ARF6 glutathionylation at Cys90 promotes ARF6 activation and contributes to erastin-induced ferroptosis in multiple cell types.

**Figure 7.**
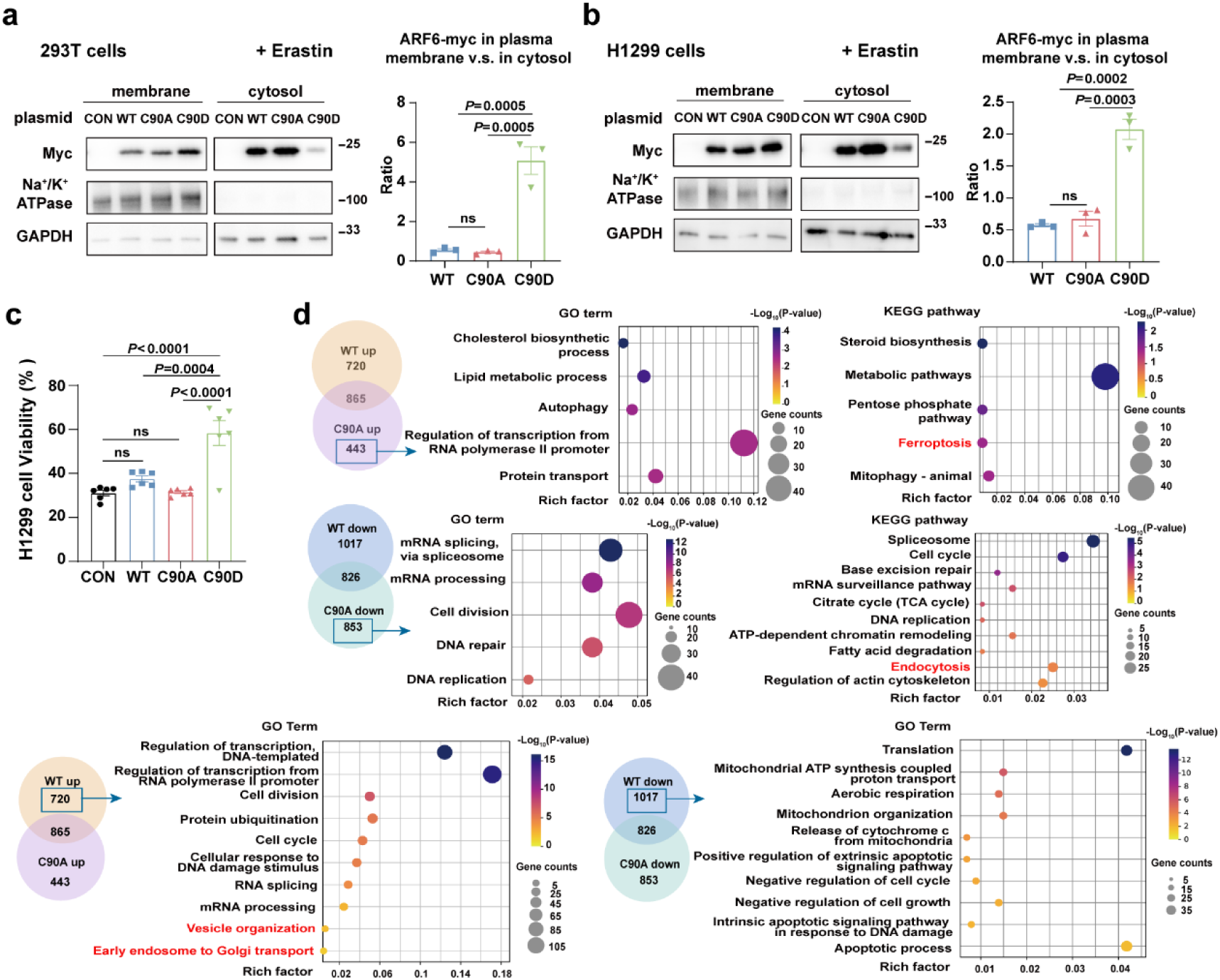
ARF6 glutathionylation at Cys90 promoted ARF6 activation and contributed to erastin-induced ferroptosis in multiple cell types. **a**: Expression of the ARF6-Myc-tag protein in 293T cells transfected with CON, WT, C90A, or C90D plasmids and treated with 10 μM erastin was analysed using western blotting. GAPDH marks the cytosol, and Na^+^/K^+^-ATPase marks the membrane. Statistical chart shows the ratio of ARF6-Myc in the plasma membrane to ARF6-Myc in the cytosol. **b**: Expression of the ARF6-Myc-tag protein in H1299 cells transfected with CON, WT, C90A, or C90D plasmids and treated with 10 μM erastin was analysed by western blotting. GAPDH marks the cytosol, and Na^+^/K^+^-ATPase marks the membrane. Statistical chart shows the ratio of ARF6-Myc in the plasma membrane to ARF6-Myc in the cytosol. **c**: The viability of H1299 cells transfected with CON, WT, C90A, or C90D plasmids and treated with 10 μM erastin was quantified using the CellTiter-Glo luminescent cell viability assay. **d**: RNA-seq analysis using ARF6-myc (WT) or ARF6-C90A-myc transfected H1299 cells, with or without erastin treatment. Venn plot, GO and KEGG enrichment for genes upregulated only in the ARF6-C90A group but not in the WT group, genes downregulated only in the ARF6-C90A group but not in the WT group, genes only upregulated in WT group but not in the ARF6-C90A group upon 10 μM erastin treatment. **e**: Schematic presentation of the effect of decreased protein S-glutathionylation caused by CHAC1-induced glutathione deprivation on cell susceptibility to ferroptosis.

## DISCUSSION

Protein-SSG levels are strongly associated with the redox state and cellular glutathione pools. Under oxidative stress, S-glutathionylation occurs spontaneously in certain proteins with available glutathione including GSH and GSSG. The present study showed a strong association among decreased glutathione pools induced by CHAC1, reduced protein-SSG levels, and ferroptosis susceptibility. By manipulating CHAC1, we found that protein S-glutathionylation conferred resistance to ferroptosis. Through glutathionylomic analysis, we identified ARF6 as a target protein that was S-glutathionylated on Cys90 to maintain the recycling of the transferrin receptor. Overall, we demonstrated that cellular glutathione deficiency induced by CHAC1 upregulation inactivated ARF6 by decreasing S-glutathionylation, thus promoting a labile iron pool and ferroptosis.

CHAC1 has been reported to be a valid biomarker for poor prognosis in cancer, including human breast and ovarian cancers, renal clear cell carcinoma, and uveal melanoma (Goebel, Berger et al., 2012, Jahn, Arvandi et al., 2017, Li, Liu et al., 2021, Liu, Li et al., 2019). CHAC1 is highly expressed in ferroptotic cells (Stockwell, 2022). Owing to its marked upregulation and occurrence in the early stages, CHAC1 is currently regarded as an indicator of ferroptosis. Similar to previous studies demonstrating that CHAC1 is one of the most upregulated genes in ferroptosis, our unbiased RNA-seq data further confirmed its specific enhancement during ferroptosis induced by erastin in H1299 cells (data not known) or APAP in primary mouse hepatocytes and livers.

The role of CHAC1 in ferroptosis has also been previously explored. CHAC1 enhances cystine-starvation-induced ferroptosis through degradation of GSH and activated GCN2-eIF2α-ATF4 pathway in human triple negative breast cancer cells (Chen, Wang et al., 2017). CHAC1 facilitates ferroptosis by accelerating GSH deprivation in retinal pigment epithelial cells with oxidative damage (Liu, Wu et al., 2023). Dihydroartemisinin effectively induces ferroptosis in primary liver cancer cells by upregulating CHAC1 expression (Wang, Li et al., 2021). Anti-CHAC1 exosomes from adipose-derived mesenchymal stem cells inhibit neuronal ferroptosis and alleviate cerebral ischemia/reperfusion injury in mice (Wang, Niu et al., 2023). CHAC1 decreased cell viability and increased the sensitivity of prostate cancer cells to docetaxel by inducing ferroptosis (He, Zhang et al., 2021). Our data showed that CHAC1 promotes erastin- and APAP-induced ferroptosis in multiple cell types. Ferroptosis is involved in APAP-induced hepatotoxicity (Lorincz et al., 2015, Niu et al., 2022, Yamada et al., 2020). Hence, the APAP-induced liver injury mouse model may be a valid in vivo model of ferroptosis. In this study, we verified the role of CHAC1 in accelerating ferroptosis in an AILI mouse model.

GSH deficiency is one of the most important events in ferroptosis; thus, the general argument in previous studies was that CHAC1 expression promotes glutathione deficiency. Our study sheds new light on the consequences of a decreased glutathione pool induced by CHAC1. With protein modification capacity, glutathione deficiency induced by CHAC1 upregulation during ferroptosis also leads to decreased protein-SSG levels, especially ARF6 S-glutathionylation. Our data show that decreased protein-SSG could also be the underlying mechanism by which glutathione deficiency promotes ferroptosis.

Oxidative stress is required for protein-SSG, which can occur either when GSSG, the GSH oxidised by oxidative stress, exchanges thiol groups with protein cysteine thiolate groups or when already oxidised protein sulfenic acid groups react with GSH(Oppong et al., 2023). Through these reactions, S-glutathionylation was observed under different oxidative conditions. Although CHAC1 is not the enzyme responsible for catalysing deglutathionylation, our study found that the upregulation of CHAC1 significantly decreased the level of protein-SSG by degrading glutathione during ferroptosis. Thus, in addition to oxidative stress, glutathione pool availability is essential for protein S-glutathionylation and subsequent ferroptosis.

Few studies have focused on the protein-SSG in APAP-induced hepatotoxicity. Yang et al. observed that glutathionylated regions were present in the liver sections of mice challenged with APAP (Yang, Greenhaw et al., 2012). Chan et al. performed a proteome-wide screening of HepaRG cells treated with APAP and characterised 898 glutathionylated peptides corresponding to 588 proteins. Glutathionylated proteins are involved in the well-known toxic effects of APAP, including energy metabolism, oxidative stress, cytosolic calcium, and mitochondrial dysfunction (Chan, Soh et al., 2018). Here, we further demonstrated aberrant protein glutathionylation upon APAP treatment in primary mouse hepatocytes, which could better preserve hepatocyte features. We identified 1105 glutathionylated peptides from 482 proteins. Consistent with the data of Chan et al. in hepatic cell lines(Chan et al., 2018), many proteins related to mitochondrial function, lipid metabolism, and energy metabolism were glutathionylated in APAP-challenged PMHs. Generally, it is recognized that protein-SSG is a protective process against oxidative stress. In this study, we demonstrated that S-glutathionylation ensures ARF6 activity for transferrin receptor recycling. Decreased glutathione pools induced by CHAC1 reduced the level of S-glutathionylation of Cys90 and inactivated ARF6, thereby promoting ferroptosis.

ARF6 is a member of the RAS superfamily. Our data suggest that ARF6 suppresses ferroptosis, consistent with previous studies (Geng & Wu, 2022, Ye et al., 2020). Pimentel et al. have reported that mechanical strain causes ROS-dependent S-glutathiolation of Ras at Cys118 and activates the Raf/Mek/Erk pathway, ultimately leading to hypertrophy in cardiac myocytes (Pimentel, Adachi et al., 2006). In the present study, we found that ARF6 glutathionylation at Cys90 decreased APAP-induced ferroptosis, which was retarded by CHAC1 overexpression. ARF6 is a Ras-related small GTPase that cycles between an active (GTP-ARF6) and inactive form (GDP-ARF6). Upon stimulation by agonists, GDP-ARF6 is activated to GTP-ARF6 by ARF6 guanine nucleotide exchange factors). Then GTP-ARF6 interacts with its effectors to regulate vesicle trafficking and is inactivated by ARF6 GTPase-activating proteins, which facilitate GTP hydrolysis to GDP (Grant & Donaldson, 2009, Maxfield & McGraw, 2004). ARF6 localises to the plasma membrane in its GTP state, and to the tubulovesicular compartment in its GDP state (Radhakrishna & Donaldson, 1997). Our data showed that the ARF6 C90A mutant, mimicking the unglutathionylated form, suppressed ARF6 expression in the plasma membrane, whereas the C90D mutant, mimicking the glutathionylated form, promoted ARF6 expression in the plasma membrane of multiple cell types. In addition, CHAC1, which catalyses glutathione degradation and reduces ARF6 S-glutathionylation at Cys90, decreases ARF6 activity in PMHs undergoing APAP-induced ferroptosis.

ARF6 plays a fundamental role in transferrin receptor (TFR) endocytosis. The TFR in the plasma membrane binds to the two Fe^3+^-bearing transferrins (TFs) and concentrates in the clathrin-coated pits. After endocytosis, the acidic pH of endosomes triggers iron release. Fe^3+^ must be reduced to Fe^2+^ by STEAP3, followed by transport across the endosomal membrane to the cytoplasm by the iron transporter, DMT1. From early endosomes, TF and TFR complexes are either delivered to the endolysosomal system for degradation or recycled directly or indirectly to the plasma membrane via the endocytic recycling compartment (ERC). Most TFRs are recycled from the ERC to the cell surface. Once TFR-TF is returned to the cell surface, iron-free transferrin is released from the TFR at a neutral extracellular pH (Anderson & Frazer, 2017, Kawabata, 2019). D’Souza-Schorey et al. reported that ARF6 promoted redistribution of TFR to the cell surface and decreased the rate of uptake of transferrin, whereas expression of *ARF6* (T27N), a dominant negative mutant, led to the intracellular distribution of TFR and inhibition of transferrin recycling to the cell surface (D’Souza-Schorey et al., 1995). In line with these data, our study found that CHAC1 inactivated ARF6 and delayed the endosomal recycling of TFR. Wang et al. reported that the loss of folliculin, a key regulator of the endocytic recycling pathway, caused a trend of reduced (not statistically significant) total iron levels but significantly increased the labile iron pool in HEK293 cells (Wang, Wu et al., 2021). Our data revealed that glutathione depletion induced by CHAC1 overexpression increases the labile iron pool during APAP-induced ferroptosis.

In conclusion, protein S-glutathionylation confers ferroptosis resistance. Upon oxidative stress, glutathione deficiency inactivates ARF6 by decreasing S-glutathionylation. Dysfunctional ARF6 delays the endosomal recycling of the TRFs and increases the labile iron pool to promote ferroptosis. This study provides novel insights into how glutathione deficiency impairs OPTMs and subsequently promotes cell ferroptosis.

## MATERIAL AND METHODS

### Animal studies

C57BL/6J mice, aged six to eight weeks, were purchased from the Experimental Animal Centre of Shanghai SLAC (Shanghai, China). *Chac1* knockout mice (*Chac1*^-/-^ mice; C57BL/6N) were constructed using CRISPR/Cas9-mediated genome engineering and were provided by Cyagen Biosciences (Suzhou, China). In this strain, exons 1–3 of the *Chac1* gene were removed, and gene expression was abolished. Male and female homozygous *Chac1*^-/-^ mice were viable and fertile. Littermate mice were used as wild-type controls (*Chac1*^+/+^). The mice were housed in a specific pathogen-free facility with 12-h light/dark cycle at 22°C in Fudan University Experimental Animal Center (Shanghai, China), were fed on a normal diet, and had ad libitum access to water. All mice were fasted overnight for approximately 12 h before APAP administration. APAP (Sigma-Aldrich, St Louis, MO, USA) was dissolved in saline at 37°C just before the experiments were conducted. The mice were injected intraperitoneally (i.p.) with saline or 300 mg/kg APAP and then euthanised to collect blood and liver samples after 6 h or at other indicated times. The protocols used in all studies were approved by the Institutional Animal Care and Use Committee of Fudan University.

### Human sample collection

The specimens consisted of nine needle biopsies obtained from patients with drug-induced liver injury (DILI) caused by nonsteroidal anti-inflammatory drugs. For healthy controls, liver tissues were obtained from the livers of transplant donors. The clinical data of the patients with DILI are shown in Supplementary Table 1, numbered from patient 1 to 9. All the patients provided informed consent to participate in this study. This study was approved by the Ethics Committee of Shanghai Jiaotong University Renji Hospital in accordance with the ethical guidelines of the 1975 Declaration of Helsinki.

### Adenovirus infection

The CHAC1 (Ad-CHAC1) and control (Ad-GFP) adenoviruses were purchased from GeneChem Technologies Co., Ltd. (Shanghai, China). Ad- CHAC1 or Ad-GFP was injected via the tail vein of each *Chac1*^-/-^ mouse at a dose of 1 × 10^8^ viral titres. After 48 h, the mice were injected i.p. with APAP (300 mg/kg) and subsequently euthanised after 6 h. For *in vitro* experiments, primary mouse hepatocytes (PMHs) were incubated with Ad-CHAC1 or Ad-GFP for 12 or 36 h, followed by treatment with APAP at the indicated time intervals.

### Small interfering RNA (siRNA)-mediated knockdown of *Chac1* and *Arf6*

We used the *Chac1* siRNA (si*Chac1*) and *Arf6* siRNA (si*Arf6*) for inhibit CHAC1 and ADP-ribosylation factor (ARF)6 expression, respectively. *SiChac1*, si*Arf6,* and corresponding negative controls (siNCs) were purchased from GenePharma (Shanghai, China). The *in vitro* transfection with si*Chac1, siArf6,* and siNC was performed using Lipofectamine 3000 (Invitrogen, USA). The target sequences of the siRNAs are listed in Supplementary Table 2.

### Cell cultures

PMHs were isolated via in situ liver perfusion with type IV collagenase (Gibco, Cat. # 17104019), following established protocols (Wang, Yang et al., 2017). The PMHs were then passed through a 70-µm cell strainer and centrifuged in 50% Percoll (Yeasen, China, Cat. # 40501ES60) at 50 × g for 5 min to separate the viable and nonviable hepatocytes. The hepatocytes were resuspended in DMEM/high glucose (VivaCell, China, Cat. # C3103-0500) supplemented with 10% FBS (ExCell Bio, China, Cat. # FSD500) and plated onto gelatin-coated dishes (Cell Biologics, Cat. # 6950).

Mouse hepatoma (Hepa1-6), murine hepatic (AML12), human embryonic kidney (293T), human colonic adenocarcinoma lung metastasis (T84), human hepatocellular carcinoma (HepG2), human oesophageal carcinoma (TE-1), human prostate cancer (DU145), human non-small cell lung cancer (H1299), human ovarian cancer (A2780), human ovarian cancer (SKOV3), and human gastric cancer (HGC27) cells were cultured in DMEM or RPMI-1640 supplemented with 10% FBS (ExCell Bio, Cat. # FSD500), 100 U/mL penicillin, 100 µg/mL streptomycin, and 0.25 µg/mL amphotericin B.

### RNA extraction and real time-quantitative polymerase chain reaction (RT-qPCR)

Total RNA was extracted using TRIzol reagent (Invitrogen, Cat. # 12183555). Subsequently, cDNA synthesis was performed using a ReverTra Ace qPCR RT Kit (TOYOBO, Cat. # QPK-201). Quantitative real-time PCR was performed using a SYBR Green PCR Kit (Yeasen, Cat. # 11203ES03). For each sample, the expression of each target gene was normalised to the expression of 36b4 or β-actin. The relative expression of genes was measured using the 2 ^(-ΔΔCt)^ method. The primer sequences are listed in Supplementary Table 3.

### Western blotting

The cells were lysed using NP40 buffer supplemented with protease and phosphatase inhibitors. Membrane and cytoplasmic proteins were isolated using membrane and cytosol protein extraction kits (Beyotime, China, Cat. # P0033). Protein levels were determined using a BCA protein assay. For western blotting, lysates were probed with specific antibodies against CHAC1 (Proteintech, Cat. # 15207-1-AP, 1:1000), GSH (Virogen, Cat. # 101-A, 1:1000), ARF6 (Proteintech, Cat. # 20225-1-AP, 1:1000), TFRC (Abcam, Cat. # ab218544, 1:4000), FLAG-tag (Sigma-Aldrich, Cat. # F1804, 1:3000), and HA-tag (Cell Signaling Technology, Cat # 3724, 1:1000). β-actin (Proteintech, Cat. # 66009-1-Ig, 1:10000), HSP90 (Proteintech, Cat. # 60318-1-Ig, 1:10000), and GAPDH (Proteintech, Cat. # 10494-1-AP, 1:5000) were used as loading control. For membrane proteins, Na^+^/K^+^ ATPase α (Santa Cruz, Cat. # sc-48345, 1:1000) was used as an internal reference. For the western blot analysis targeting GSH, the sample preparation did not include the addition of reducing agent β-mercaptoethanol, and total proteins stained with Ponceau S were used as the control. Goat anti-rabbit IgG HRP (Affinity, Cat. # S0001, 1:3000) and goat anti-mouse IgG HRP (Affinity, Cat. # S0002, 1:3000) were used as secondary antibodies. Proteins were visualised using an ECL kit (Cat. # 180-501) and a Tanon-4200 gel imaging system.

### Arf6 pulldown activation assay

The Arf6 pulldown activation assay was conducted using a commercial kit (New EastBiosciences, China, Cat. # 82401). Briefly, cells were lysed in NP-40 lysis buffer (Beyotime, Cat. # P0013F) containing 1% Phenylmethanesulfonyl fluoride and centrifuged to collect the supernatant. The cell lysate containing 2 mg total protein was adjusted to 1 mL with lysis buffer and supplemented with 2 μg of active Arf6 monoclonal antibody and 30 μL of protein A/G agarose. The mixture was rotated for 4 h at 4°C. GTPγS-treated protein was used as a positive control. The agarose beads were then washed five times with buffer, with each wash lasting 5 min, followed by centrifugation at 400 × g for 1 min to remove the supernatant. After the final wash, all supernatants were removed and the sample was resuspended in an equal volume of 2× SDS-PAGE loading buffer. The samples were boiled for 5 min and subjected to western blotting for visualisation and analysis of the pulled-down samples.

### Biochemical assays and histopathological analyses

Serum alanine aminotransferase (ALT) and aspartate aminotransferase (AST) levels were measured using commercial assay kits (Nanjing Jiancheng, Cat. # C009-2-1, C010-2-1) according to the manufacturer’s instructions. Formalin-fixed and paraffin-embedded liver tissue sections were stained with haematoxylin and eosin (H&E) and analysed. DNA damage was detected using a terminal dUTP nick-end labelling (TUNEL) assay kit (Roche, Cat. # 11684817910). Malondialdehyde (MDA) levels were measured using the appropriate assay kit (Dojindo, Japan, Cat. # M496) and normalised to the protein concentration according to the manufacturer’s instructions.

### Immunohistochemistry

Immunohistochemistry was performed on formalin-fixed paraffin-embedded liver tissue sections. Sections were incubated with an anti-rabbit CHAC1 (Proteintech, Cat. # 15207-1-AP) and anti-rabbit 4HNE (Abcam, Cat. # ab46545) antibody at 4°C overnight. Subsequently, tissues were incubated with a peroxidase-conjugated rabbit anti-goat secondary antibody at room temperature for 1 h. Tissue sections were counterstained with DAB and haematoxylin. IHC staining for CHAC1 and 4-hydroxynonenal (4-HNE) was analyzed using histological scoring. The scoring criteria were as follows: score =0, no staining; score =1, 1–25%; score =2, 26–50%; score =3, 51–75%; and score =4, 76–100%. The intensity scores represented the average value: 0 (none), 1 (weak), 2 (moderate), and 3 (strong). Each sample was evaluated in a blinded manner by a senior pathologist and two researchers. The final score for each sample was the average of three scores from the researchers. The quantity and intensity scores were then multiplied to obtain a total score ranging from 0 to 12, representing the histological score.

### Immunofluorescence staining

Subcellular localisation and protein expression were assessed using immunofluorescence staining. PMHs were seeded onto slides and infected with AdGFP or AdCHAC1 adenoviruses for 36 h, followed by a 6-h treatment with 20 mM APAP. Slides were incubated overnight with an anti-mouse HA (Santa Cruz, Cat. # sc-7392, 1:200) and anti-rabbit TFRC (Abcam, Cat. # ab214039, 1:200), followed by incubation with a fluorescent secondary antibody at room temperature for 1 h. DAPI was used for nuclear staining and the cells were incubated for 10 min. Fluorescent images were captured using a confocal microscope (Leica).

### GSH and GSSG concentration determination

GSH and GSSG levels were quantified using a GSSG/GSH Quantification Kit (Dojindo, Cat. # G263) according to the manufacturer’s instructions. In addition, GSH and GSSG levels were normalised to the protein concentration.

### Cell viability

The cells were seeded in opaque 96-well microplates and exposed to the specified treatments. Cell viability was assessed using the CellTiter-Glo Luminescent Cell Viability Assay Kit (Promega, Cat. # G7570) according to the manufacturer’s protocol. Luminescence intensity from each well was measured with the Synergy™ H1 Hybrid Multi-Mode Microplate Reader (BioTek, USA). Cell viability was expressed as a percentage of control cells.

Calcein-AM/PI Double Staining Kit was also used to assess cell viability. Cells were washed twice with PBS before staining and then incubated with staining solution at 37°C for 30 min, following the manufacturer’s protocol (Dojindo,, Cat. #C542). After three washes with PBS, the cells were imaged using a fluorescence microscope (Nexcope, China), and ImageJ software (version 1.53c) was used to count fluorescent cells.

### Lipid peroxides determination

Cellular lipid peroxidation levels were determined using a C11 BODIPY 581/591 fluorescent probe (Dojindo, Cat. # L267) according to the manufacturer’s instructions.

### Measurement of Fe^2+^ level

FerroOrange (Dojindo, Cat. # F374) was used to assess the Fe^2+^ levels. PMHs were seeded in 12-well plates at a density of 1 × 10^5^ cells/well. After cell adhesion, the cells were transfected with *Arf6* siRNA for 48 h or infected with AdGFP or AdCHAC1 adenovirus for 36 h, and then treated with 20 mM APAP for 6 h. Following the removal of the previous culture medium, the cells were washed twice with PBS. Subsequently, a working solution of FerroOrange fluorescent probe at a concentration of 1 μmol/L was added to the cells. The cells were incubated at 37°C in the dark for 30 min, and then incubated with DAPI for 10 min for nuclear staining. Finally, the cells were observed and images were acquired using a fluorescence microscope (Nexcope). ImageJ software was used to quantify the fluorescence intensity of the FerroOrange.

### Measurement of transferrin

The transferrin levels were measured using a transferrin probe (Invitrogen)Cat. # T23364), according to the manufacturer’s instructions. In addition, PMHs were seeded in 12-well plates at a density of 1 × 10^5^ cells/well. After cell adhesion, the cells were infected with AdGFP / AdCHAC1 adenovirus for 36 h, and then treated with 20 mM APAP for 6 h. Subsequently, the cells were placed on ice for 10 min, washed twice with pre-chilled DMEM containing 1% BSA, and then incubated in the dark with a final concentration of 25 μg/mL probe dye at 37°C for 30 min. After incubation, the cells were washed twice with pre-chilled DMEM containing 1% BSA. The cells were fixed and incubated with Hoechst stain for 10 min for nuclear staining. Fluorescence images were captured using a confocal microscope (Leica), and transferrin levels were quantified using ImageJ software.

### Molecular docking

ARF6 protein sequences were obtained from the UniProt database using ID P62331. The total length of the sequence was 175 amino acids, and there was no protein crystal structure with a full-length sequence. Therefore, the structure of ARF6 was predicted using Alphafold 2 (https://alphafold.ebi.ac.uk/) and selected for subsequent protein docking and structural modification.

The MOE 2015.10 (CCG, Ottawa, Canada) program was used to assess the predicted structure, structure preparation, pocket search, and docking. The reliability of the protein models was validated using Ramachandran plots. A total of 23 binding sites were searched by MOE, which ranked at the top, and were used for further studies. The amino acid site ID are Leu21, Asp22, Ala23, Ala24, Gly25, Lys26, Thr27, Thr28, Thr41, Ile42, Pro43, Thr44, Asp63, Val64, Gly65, Gly66, Gln67, Asn122, Lys123, Asp125, Cys155, Ala156, and Thr157. The ligand structures of GTP and GDP were downloaded from PubChem (https://pubchem.ncbi.nlm.nih.gov/). The ligand was docked to ARF6 using the induced fit-docking method. To investigate the influence of GSS modification on the binding of the ligand to ARF6, amino acid residue cysteine residue 90 (Cys90) was selected as the modification site. The modified structure was prepared using MOE through energy minimisation, optimised by molecular dynamics with an amber99sb forcefield and 500 ps simulation. The representation was prepared using PyMOL 2.4.0.

### RNA sequencing (RNA-seq) and data analysis

RNA purified from the cultured cells was converted into cDNA libraries and assessed for quality using an Agilent 2100 Bioanalyzer. Sequencing was performed using a BGI DNBSEQ-T7 sequencer (OE Biotech Co., Ltd., (Shanghai, China). Raw RNA sequencing data were deposited in the NCBI SRA database under the accession number PRJNA1086892. Differentially expressed genes (DEGs) were identified using Cuffdiff and defined as *P* < 0.05. Gene Ontology (GO) and Kyoto Encyclopedia of Genes and Genomes (KEGG) enrichment analyses of DEGs was performed using the DAVID online analysis system (https://david.ncifcrf. gov/homejsp).

### Enrichment of protein-SSG and LC–MS/MS

PMHs in 10-cm dishes treated as indicated were scraped using cell scrapers and pelleted by centrifugation. The cell pellets were heated at 95℃ for 10 mins, and then lysed in four times the volume of lysis buffer (1% SDS, 1% protease inhibitor cocktail, and 25 mM iodoacetamide) under ultrasonication on ice. Iodoacetamide (25 mM) was added, and the alkylation reaction was performed at room temperature in the dark for 1 h. The samples were centrifuged at 4°C for 10 min at 12,000 × g to remove the cell debris, and the supernatant was transferred to a new centrifuge tube for protein concentration determination using a BCA kit.

For the reduction reaction, 480 µg of proteins (1 µg/µL) were treated in 25 mM HEPES (pH 7.5) with 1 M urea, 2.5 µg/mL Glrx 1M, 0.25 mM GSSG, 1 mM NADPH, and 4 U/mL Glutathione Reductase, and incubated at 37°C for 10 min, immediately placed on ice, and transferred to a 0.5-mL Amicon Ultra 10K filter. Excess reagents were removed by buffer exchanged with 3× 8 M urea (pH 7.0) resulting in a final volume of 30–40 μL. Next, each protein channel was labelled with its respective iodo-TMT reagent and incubated for 1 h at room temperature in the dark. DTT (final concentration: 20mM) was added and the cells were incubated for another 15 min in the dark.

Proteins were precipitated with acetone, centrifuged, washed, and resuspended in 200 mM triethylammonium bicarbonate buffer for overnight digestion with trypsin at a 1:50 enzyme-to-substrate ratio. Digested samples were reduced at 37°C for 60 min using 5 mM dithiothreitol and subsequently alkylated in the dark at room temperature with 11 mM iodoacetamide for 45 min. The peptides were purified using Strata X SPE columns and vacuum-dried before resuspension in 1X TBS. The samples were incubated overnight at 4°C with anti-TMT resin in an end-to-end mixture. Following incubation, the supernatant was removed and the resin was washed eight times with one column volume of TBS (5 min per wash), followed by three washes with one column volume of water. The components were then eluted with four column volumes of TMT elution buffer (50% acetonitrile and 0.4% trifluoroacetic acid), and the eluates were dried under vacuum. The peptides were desalted using C18 ZipTips prior to LC-MS/MS analysis.

MS/MS analysis was conducted using MaxQuant software (version 1.6.15.0) to search 17,132 sequences, incorporating a reverse decoy database and contaminant database to manage the false discovery rate (FDR) and contamination issues. The settings included trypsin/P enzyme allowing up to two missed cleavages, peptide criteria with a minimum length of seven amino acids and a maximum of five modifications, mass tolerances of 20 ppm for precursor ions in the first search and 4.5 ppm in the main search, and 20 ppm for fragment ions. The quantification method was set to iodoTMT-6plex, with both protein and PSM identification FDRs set to 1%, requiring at least one unique peptide for protein identification. IodoTMT-based proteomics and analyses were performed by Jingjie PTM Biolab Co., Ltd. (Hangzhou, China).

### Statistical analysis

Data were presented as mean ± SEM. GraphPad Prism software (version 9.0; San Diego, CA, USA) was used for the calculations, statistical analyses, and graphics generation. Two-parameter comparisons were performed using a two-tailed Student’s t-test. Multiple-group analyses were performed using one-way analysis of variance (ANOVA), followed by the recommended post hoc tests using GraphPad Prism software (version 9.0). To compare kinetic differences, a two-way ANOVA test was used. Statistical significance was set at *P* < 0.05.

## ACKNOWLEDGEMENTS

This work was supported by the National Natural Science Foundation of China [(NSFC 82370597 (X.L.), NSFC 32171175 (X.L.), and NSFC 82270619(Y.M.)] and the National Key R&D Program of China [2022YFC3502101(Y.M.)]. We apologise to those whose work was relevant to, but not cited in, this study due to limited space.

## AUTHOR CONTRIBUTIONS

Conceptualisation, X.L., Y.J., and Y.M.; Methodology, X.T. and Z.L.; Formal Analysis, X.L. and Y.J.; Investigation, Y.J., Y.Z., Y.Q., T.Y., B.N., X.L., L.Y., Y.W., Y.Z., Y.D., and Q.X.; Writing-Original Draft, Y.J. and X.L.; Writing-Review and Editing, Y.Z. and X.L.; Funding Acquisition, X.W., Y.M., and X.L.; Resources, Q.X., X.W., and X.L.; Visualisation, Y.J., Y.Z., and X.L.

## INCLUSION AND DIVERSITY

We support inclusive, diverse, and equitable conduct of research.

## RESOURCE AVAILABILITY

### Lead contact

Further information and requests for resources and reagents should be directed to and will be fulfilled by the lead contact, Xiaobo Li.

### Materials availability

This study did not generate new unique reagents

### Data and code availability

- Mass spectrometry proteomic data were deposited in the ProteomeXchange Consortium (https://proteomecentral.proteomexchange.org) via the iProX partner repository with the dataset identifier PXD050569.
- Raw RNA-Seq data were deposited in GEO and are publicly available as of the date of publication under the accession number PRJNA1086892.
- This paper does not report original code.
- Any additional information required to reanalyse the data reported in this paper is available from the lead contact upon request.

## Supplementary figures

**Supplementary Figure 1, related to Figure 1.**
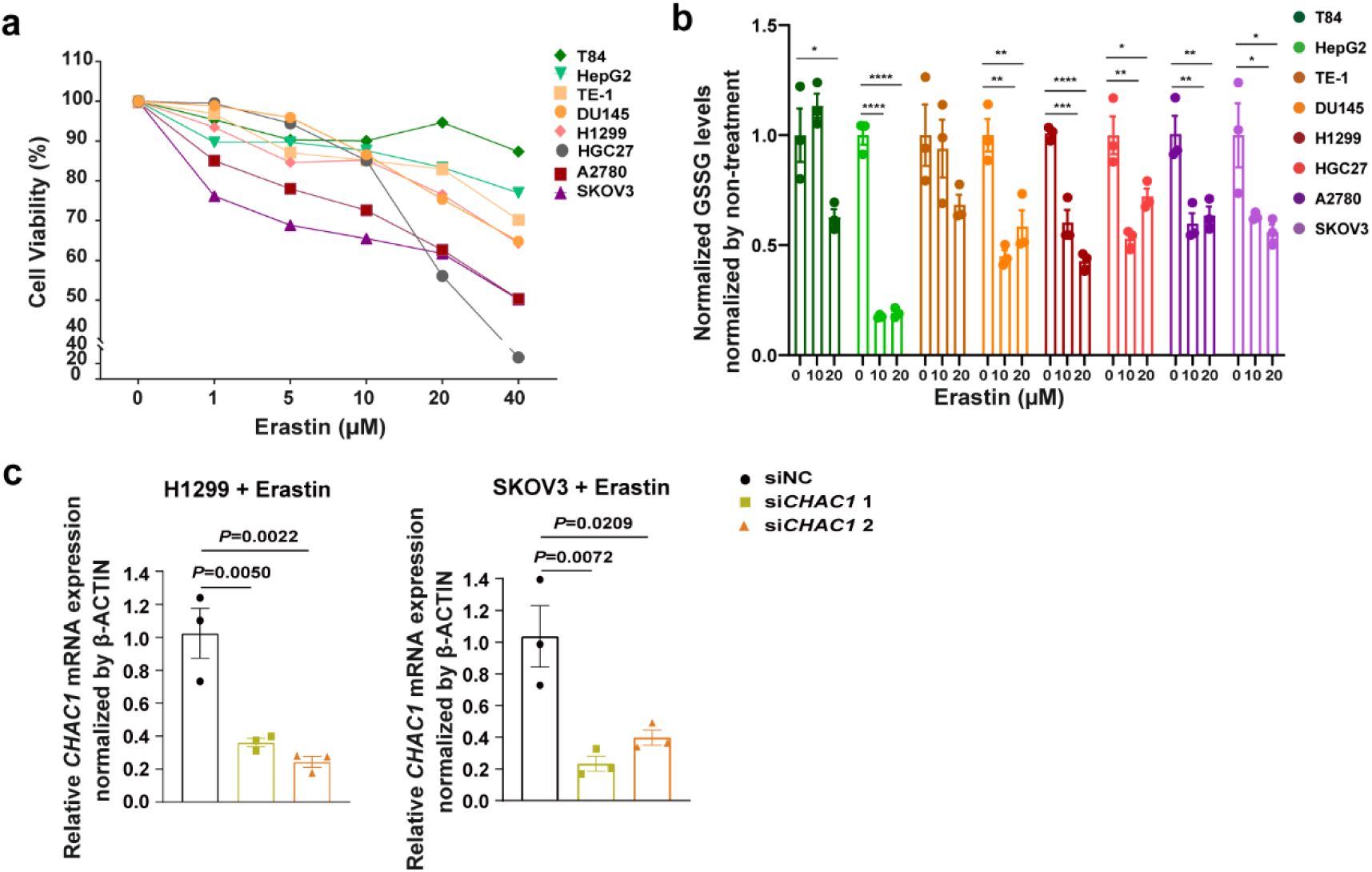
**a**: Cell viability of eight human cell lines, treated with DMSO or 1, 5, 10, 20, and 40 μM erastin, was measured by a CellTiter-Glo luminescent cell viability assay. **b**: Quantification of GSSG in eight human cell lines treated with DMSO, or 10 μM or 20 μM erastin. c: H1299 and SKOV3 cells transfected with *CHAC1* siRNA and then treated with 10 μM erastin. The mRNA level of *CHAC1* was analysed by RT-qPCR.

**Supplementary Figure 2, relate to Figure 2.**
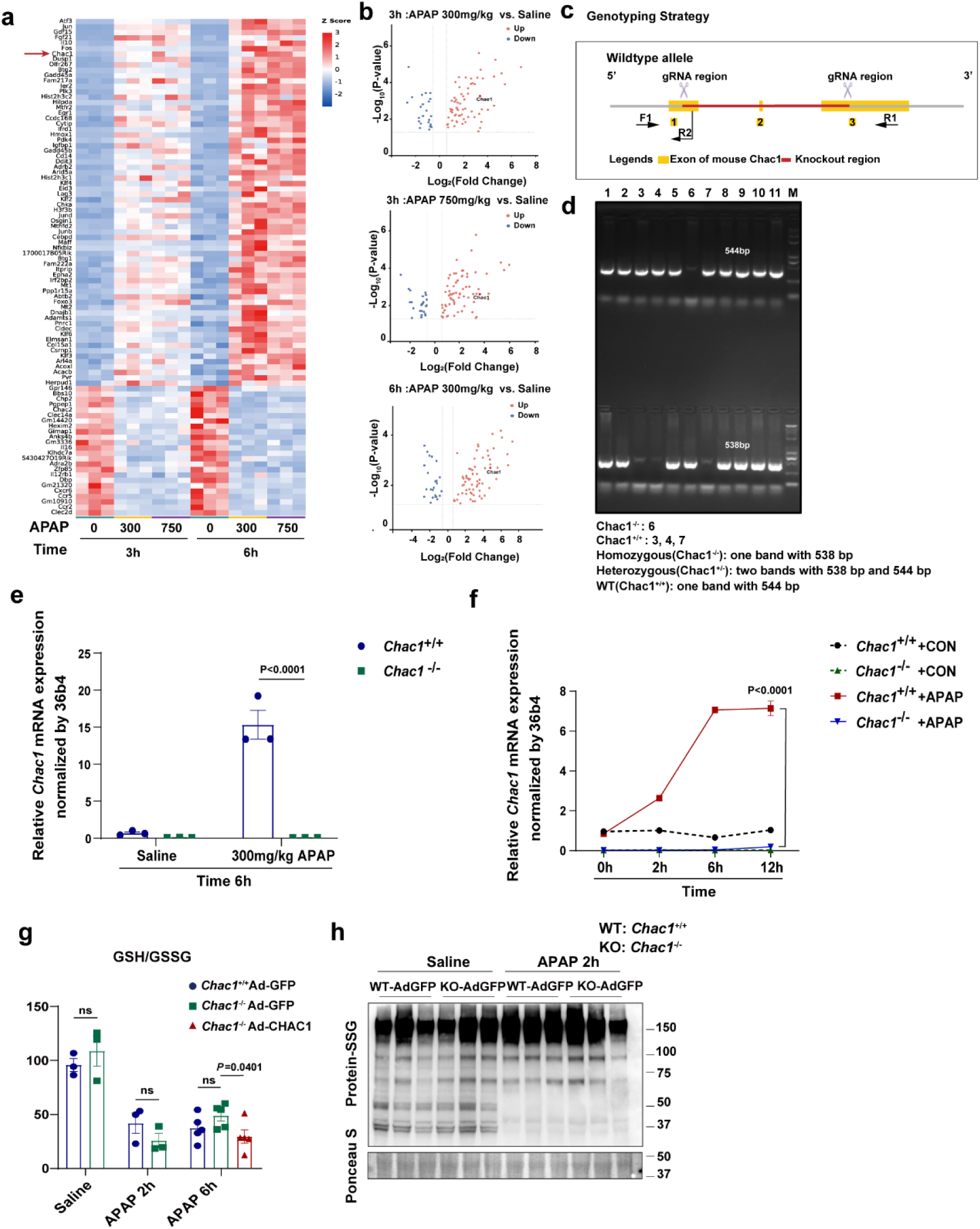
**a**: The heatmap shows the relative levels of upregulated and downregulated genes in the liver tissue of mice treated with 300 mg/kg APAP for 3 h, 750 mg/kg APAP for 3 h, 300 mg/kg APAP for 6 h, and 750 mg/kg APAP for 6 h compared to the saline group. The heatmap was ranked by fold change of 750 mg/kg APAP for 3 h group to saline for 3 h group. (Fold change ≥ 1.5, *P* < 0.05) (*n* = 3 mice/group). **b**: Volcano plots shows upregulated and downregulated genes from RNA transcriptome data of groups treated with 300 mg/kg APAP for 3 h, 750 mg/kg APAP for 3 h, and 300 mg/kg APAP for 6 h compared to the saline group. (Fold change ≥ 1.5, *P* < 0.05) (n = 3 mice/group). **c**: Strategies for *Chac1* knockout and gene identification. The *Chac1* gene (NM_026929) has three exons, with the ATG start codon in exon 1 and the TGA stop codon in exon 3. Exons 1–3 were selected as the target sites. Cas9 and gRNA were co-injected into fertilized eggs to produce for *Chac1* knockout (KO) mice. **d**: Genotyping of experimental mice. **e**: *Chac1* mRNA expression in liver tissues from *Chac1^+^*^/+^ or *Chac1^-^*^/-^ mice treated with saline or 300 mg/kg APAP for 6 h analysed by RT-qPCR (*n* = 3 mice/group, *t* test). **f**. *Chac1* mRNA expression in PMHs from *Chac1^+^*^/+^ or *Chac1^-^*^/-^ mice treated with DMEM or 20 mM APAP for 2 h, 6 h, and 12 h analysed by RT-qPCR. **g**: The ratio of GSH/GSSG in liver tissues of *Chac1^+^*^/+^ Ad-GFP, *Chac1^-^*^/-^ Ad-GFP and *Chac1^-^*^/-^ Ad-CHAC1 mice treated with saline or 300 mg/kg APAP for 2 h and 6 h. **h**: S-glutathionylated proteins in liver tissues of *Chac1^+^*^/+^ Ad-GFP and *Chac1^-^*^/-^ Ad-GFP mice treated with saline or 300 mg/kg APAP for 2 h. Ponceau S staining was used as internal reference. APAP, acetaminophen; PMH, primary mouse hepatocyte

**Supplementary Figure 3, related to Figure 3.**
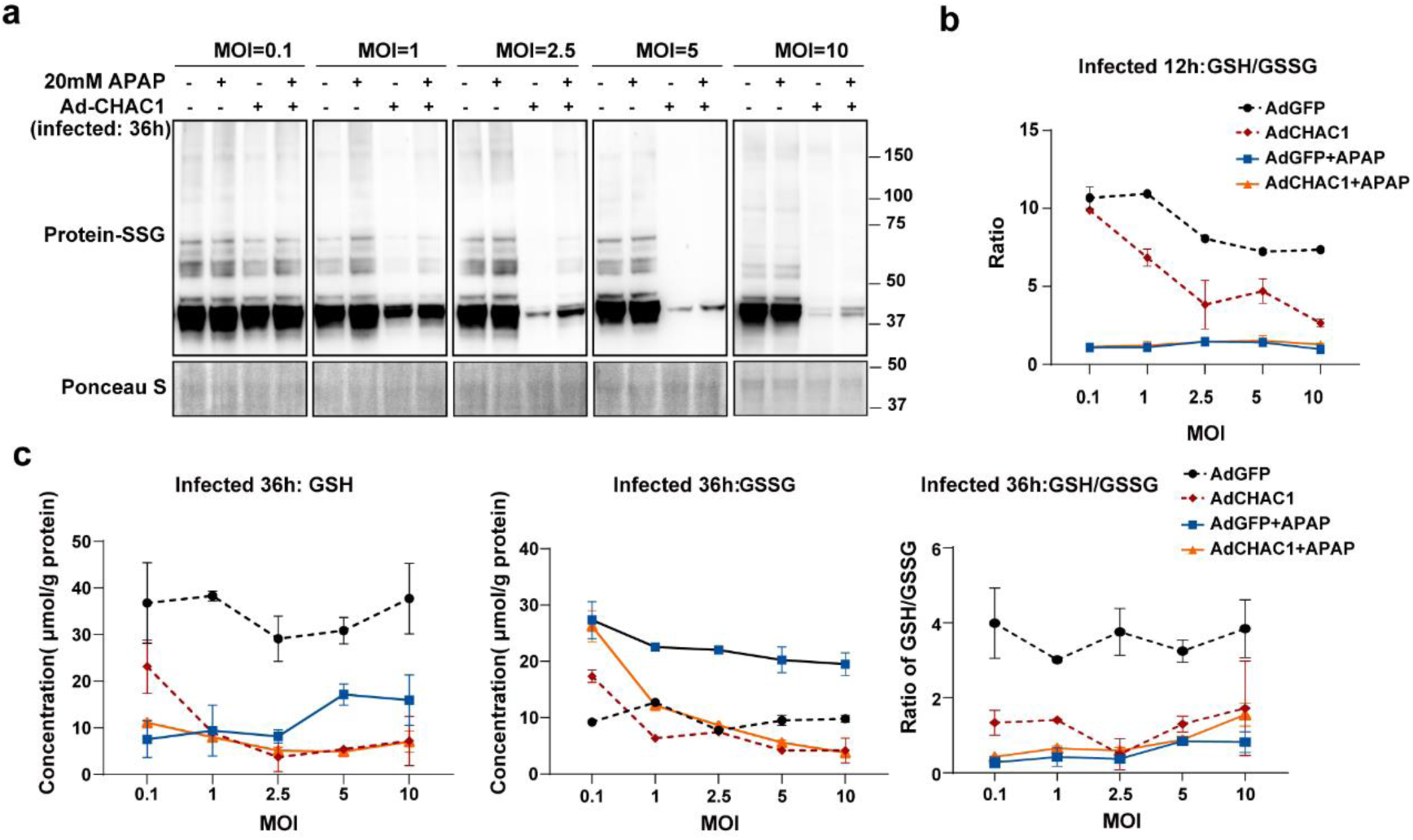
**a**: S-glutathionylated proteins in PMHs infected with Ad-GFP or Ad-CHAC1 adenovirus with MOI=0.1, 2.5, 5, and 10 for 36 h and then treated with 20 mM APAP for 6 h were analysed by western blotting with anti-glutathione. Ponceau S staining was used as internal reference. **b**: The ratio of GSH/GSSG in PMHs infected with Ad-GFP or Ad-CHAC1 adenovirus with MOI=0.1, 2.5, 5, 10 for 12 h and then treated with 20 mM APAP for 6 h. **c**: Quantification of GSH and GSSG and the ratio of GSH/GSSG in PMHs infected with Ad-GFP or Ad-CHAC1 with MOI=0.1, 2.5, 5, and 10 for 36 h and then treated with 20 mM APAP for 6 h. APAP, acetaminophen; GSH, glutathione; GSSG, oxidised glutathione

**Supplementary Figure 4, related to Figure 3.**
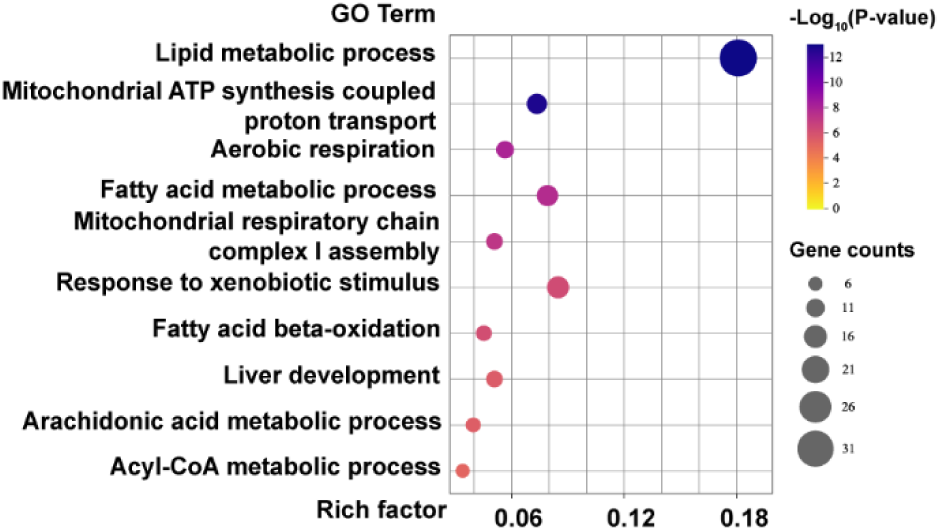
GO enrichment of differentially modified glutathionylated proteins both under APAP stimulation and CHAC1 overexpression.

**Supplementary Figure 5, related to Figure 5.**
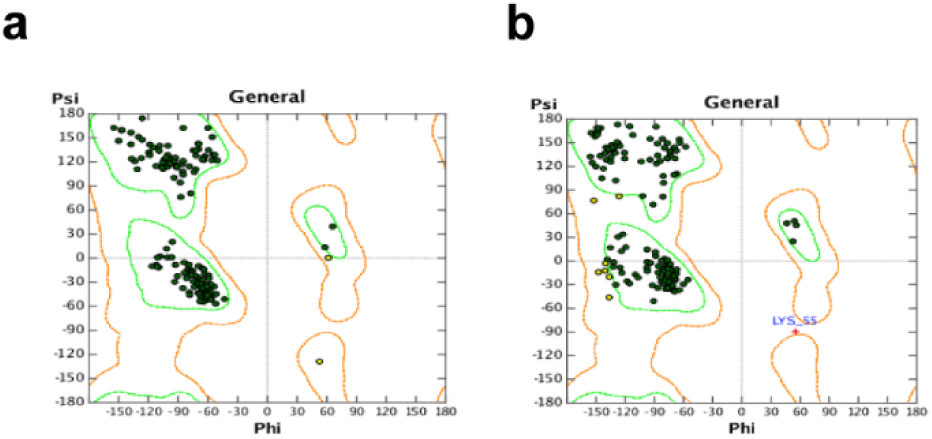
**a**: Ramachandran diagram was used to evaluate the rationality of the ARF6 protein structure. Abscissa Phi (φ) and ordinate Psi (ψ) were used to define the rationality of the geometric structure of amino acids. **b**: Ramachandran was used to evaluate the rationality of the Cys90 glutathionylated ARF6 protein structure. Abscissa Phi (φ) and ordinate Psi (ψ) were used to define the rationality of the geometric structure of amino acids.

**Supplementary Figure 6, related to Figure 6.**
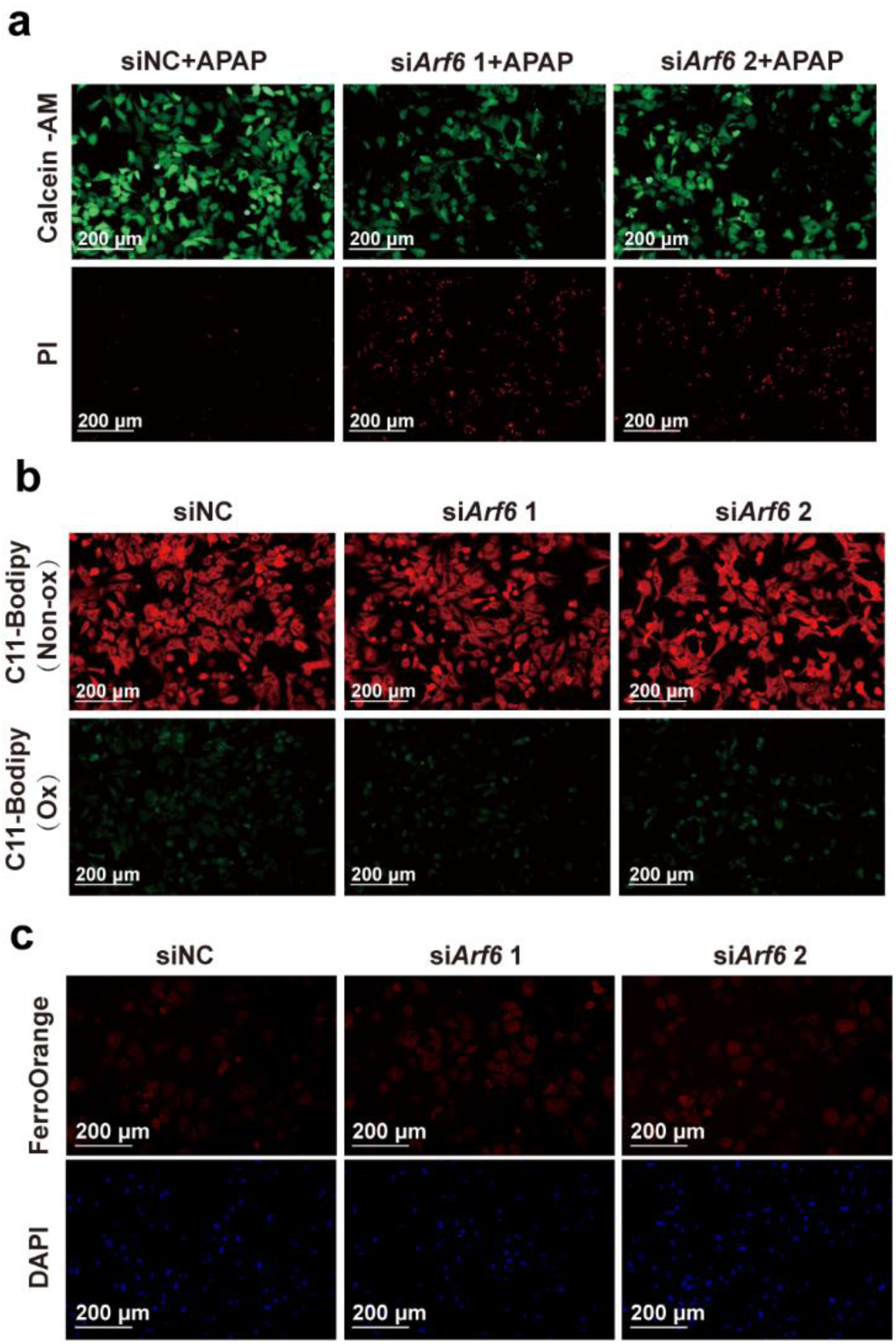
**a**: PMHs were transfected with *Arf6* siRNA and then treated with 20 mM APAP. Representative images of calcein-AM/PI live/dead cell staining (Green: live cells; red: dead cells, Scale bars = 200 μm). **b**: Representative images of C11 Bodipy 581/591 fluorescent probe of PMHs transfected with *Arf6* siRNA (Red: non-lipid oxidation; green: lipid oxidation, Scale bars = 200 μm). **c**: Representative images of FerroOrange fluorescent probe of PMHs transfected with *Arf6* siRNA (Red: FerroOrange; blue: DAPI, Scale bars = 200 μm). APAP, acetaminophen; PMH, primary mouse hepatocyte

**Supplementary Table 1.** Clinical and biological data of the patients with drug-induced liver injury.

| Patient | Sex/age | Clinical type | Drug | Drug type | Serum ALT (U/L) | Serum AST (U/L) | Serum ALP (U/L) | Serum TBIL (μmol/L) | Serum DBIL (μmol/L) |
| --- | --- | --- | --- | --- | --- | --- | --- | --- | --- |
| 1 | F/54 | Cholestatic | Acetaminophen | NSAID | 173 | 112 | 316 | ND | ND |
| 2 | F/49 | Cholestatic | Acetaminophen | NSAID | 484 | 635 | 244 | 66.2 | 21.2 |
| 3 | F/33 | Mixed | Acetaminophen | NSAID | 598 | 164 | 213 | ND | ND |
| 4 | M/65 | Cholestatic | Ibuprofen | NSAID | 152 | 54 | 194 | 63.7 | 50.2 |
| 5 | M/26 | Mixed | Not specified | NSAID | 435 | 313 | 435 | 36.81 | 23.94 |
| 6 | M/54 | Hepatocellular | Not specified | NSAID | 774 | 897 | 253 | 242 | 143.5 |
| 7 | M/32 | Hepatocellular | Not specified | NSAID | 526 | 229 | 87 | 22 | 7.9 |
| 8 | F/28 | Hepatocellular | Not specified | NSAID | 1560 | 1026 | 193 | 109.5 | 75.5 |
| 9 | F/69 | Hepatocellular | Compound Paracetamol and Amantadine Hydrochloride Capsules | NSAID | 993 | 730 | 234 | 30.6 | 23.2 |

**Supplementary Table 2.**
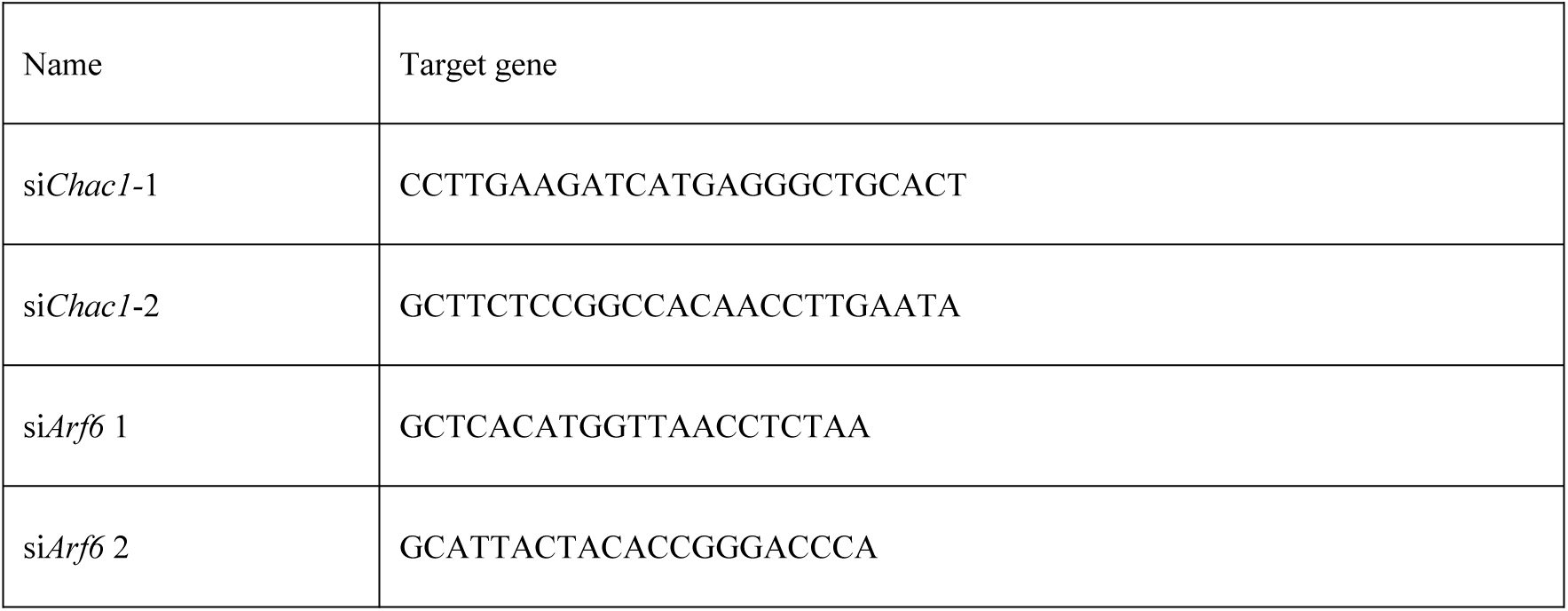
The target sequences of siRNA.

**Supplementary Table 3:**
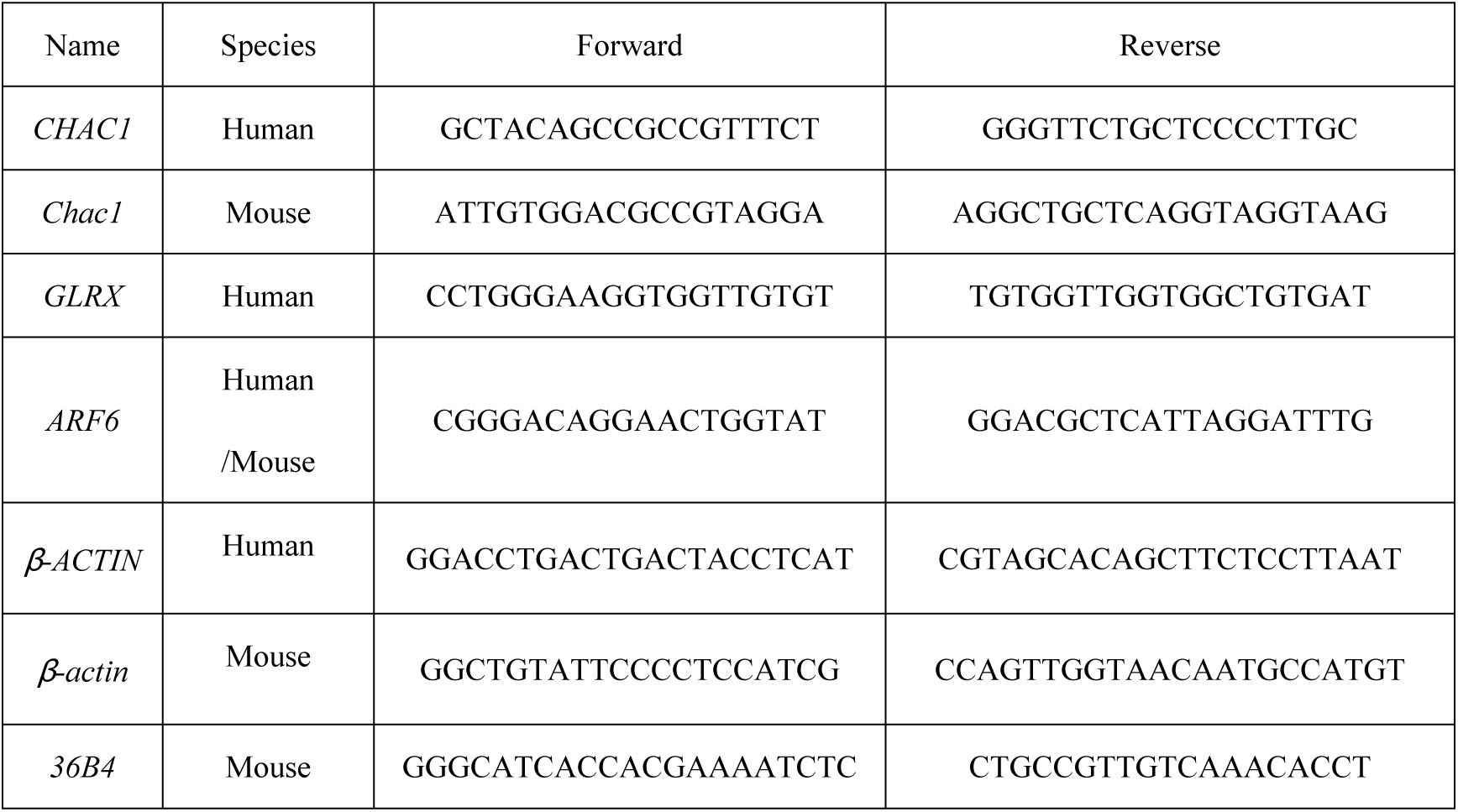
Primers used in RT-qPCR analysis.

